# Integration of Proxy Intermediate Omics traits into a Nonlinear Two-Step model for accurate phenotypic prediction

**DOI:** 10.1101/2025.03.14.643213

**Authors:** Hayato Yoshioka, Tristan Mary-Huard, Julie Aubert, Yusuke Toda, Yoshihiro Ohmori, Yuji Yamasaki, Hisashi Tsujimoto, Hirokazu Takahashi, Mikio Nakazono, Hideki Takanashi, Toru Fujiwara, Mai Tsuda, Akito Kaga, Jun Inaba, Yushiro Fuji, Masami Yokota Hirai, Yui Nose, Kie Kumaishi, Erika Usui, Shungo Kobori, Takumi Sato, Megumi Narukawa, Yasunori Ichihashi, Hiroyoshi Iwata

## Abstract

Intermediate omics traits, which mediate the effects of genetic variation on phenotypic traits, are increasingly recognised as valuable components of genetic evaluation. In particular, rhizosphere microbiota play a crucial role in plant health and productivity; however, their complex interactions with host genetics remain challenging to model. Although two-step modeling frameworks have been proposed to integrate intermediate omics traits into phenotype prediction, existing approaches do not incorporate nonlinear relationships between different omics layers. To address this, we have proposed a two-step phenotype prediction framework that integrates genomic, rhizosphere microbiome, and metabolome (meta-metabolome) data, while explicitly capturing omics-omics nonlinearities. The first step is to predict meta-metabolome traits from genetic and microbial features, thus effectively isolating them from the environmental noise. In this process, intermediate “proxy” omics traits are generated as general biological information to provide robust models. The second step utilises this “proxy” to enhance the accuracy of the phenotype prediction. We compared the linear model (Best Linear Unbiased Prediction, BLUP) and the nonlinear model (Random Forest, RF) at each step, as demonstrated through simulations and empirical analysis of a multi-omics soybean dataset in which nonlinear modeling captures intricate omics interactions. Notably, our approach enables phenotype prediction without requiring the original meta-metabolome data used in model training, thereby reducing reliance on costly omics measurements. This framework integrates intermediate omics traits into genomic prediction to improve prediction accuracy and provide solutions for deeper insights into plant-microbiome interactions.

## 1 Introduction

In breeding and genetics, intermediate omics traits, which mediate the effects of genetics on phenotypes of interest, have attracted attention for comprehensive genetic analysis. In particular, rhizosphere microorganisms play a crucial role in plant health by protecting plants against pathogens, improving nutrient uptake, and also enhancing stress resilience (Kristin & Miranda, 2013; Alemu et al., 2024). Investigating rhizosphere microbiota using multi-omics data provides an opportunity to uncover the genetic basis of plant-microorganism symbiosis. This knowledge is key to developing breeding and cultivation strategies that harness the beneficial properties of microbiota and their metabolites (Marco et al., 2022).

Genomic Best Linear Unbiased Prediction (GBLUP), a method widely used in plant and animal breeding, predicts breeding values using genome-wide DNA markers (Meuwissen et al., 2001). This model is defined as follows:

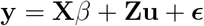

where **y** is a vector of phenotypes, **X** is a design matrix for fixed effects, *β* is a vector of fixed effects, *Z* is a design matrix linking phenotypic records to genetic values, **b** is a vector of additive genetic effects for an individual, 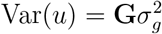 with **G** as the genomic relationship matrix (GRM), and 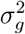 is the genetic variance, and *ϵ* is a vector of random normal deviates with variance 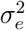.

One approach for incorporating omics into genetic evaluations is to include omics-derived traits as correlated variables in a multi-trait model (Hayes et al., 2017). This model allows for the estimation of breeding values (EBVs) for all individuals, accommodates incomplete omics data, and extends traditional models for genetic evaluation. However, because the model uses all omics-data sets as the inputs for the modeling, the prediction equations require periodic updates. Weishaar et al. (2020) proposed a theoretical framework for breeding value by incorporating microbiota composition as an intermediate trait of feed efficiency. This model uses a selection matrix to partition the breeding value into contributions from genes that affect the microbiota and those that directly influence the target traits.

Mancuso et al. (2019) proposed a hierarchical model for human genetic studies that integrates intermediate omics traits into phenotype predictions. Their model consists of two equations: First, the phenotype (**y**) is modeled as a linear function of the expression of the genes of interest, residual genetic effects, and environmental effects.

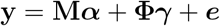

where **M** is a matrix of gene expression levels, ***α*** is a vector of associated effect sizes, **Ф** is the centered and variance-standardized genome-wide genotype matrix at each SNP, γ represents the pleiotropic effects of **Ф** on **y**, and ***e*** represents environmental effects, which follow 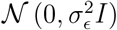. The expression levels (**M**) were modeled as a linear function of genotypes, weighted by the expression quantitative trait loci (eQTL) effects, given by **M** = **ФW**, where **W** is the eQTL effect-size matrix. This two-step approach effectively models the genetic effects on the phenotype into one that directly affects the phenotype and one that affects the phenotype through intermediate effects.

Christensen et al. (2021) extend this two-step framework by combining two linear equations. These models provide genetic evaluations that separate direct genetic effects from those mediated by omics traits. However, the assumption of linearity may limit the ability to capture any complex interactions within omics data and reduce prediction accuracy.

Recently, Zhao et al. (2022) introduced a neural network model to predict phenotypes based on intermediate omics traits. Although nonlinear approaches have been shown to be effective in predicting phenotypes, the nonlinearity in these approaches is limited to the prediction task from omics to phenotype and has not been applied to the prediction task from omics to omics.

In this study, we extended the two-step framework to extract nonlinear relationships between omics. The application of a nonlinear two-step model to eliminate environmental factors inherent in omics-omics is unprecedented. This nonlinear model can be effectively used to separate genetic traits from environmental factors, particularly in complex omics-omics relationships.

Therefore, we have proposed a novel approach to extract relevant gene and microbial information from meta-metabolome as intermediate omics traits. In the first step, intermediate omics traits are predicted using linear or nonlinear functions of the explanatory variables, effectively isolating them from environmental noise. In the second step, these predicted traits serve as “proxies” to enhance phenotype prediction. In each step, we compared the linear and nonlinear modeling. To achieve this, we employed BLUP for the linear model and Random Forest RF (Breiman, 2001) for the nonlinear model. RF is a robust machine learning method widely used in genomic science (Qi, 2012; Chen et al., 2012). RF was supposed to capture complex nonlinear patterns, which led to the leverage of intricate omics data and improved the predictive performance.

As a highlight of our model, we generate a predicted “proxy” for intermediate omics traits, allowing for model enhancement. The “proxy” is sorely based on genetic and microbiome information, making the model robust for other predictions in different environments. Once established, the model can be applied simultaneously, without requiring the original meta-metabolome data used during training. Thus, it reduces the reliance on costly omics data, such as metabolome, every time for the modeling.

This article presents a machine learning-based two-step phenotype prediction model for multi-omics. First, we present methods for incorporating linear and nonlinear models. After presenting the methods, we performed simulations to show how the nonlinear model can extract information from intermediate omics traits in linear and nonlinear settings. Subsequently, we applied the model to our multi-omics soybean dataset to demonstrate its potential to improve phenotype prediction.

## 2 Materials and Methods

### Soybean multi-omics data

We analyzed the multi-omics data obtained from the same *N* = 198 soybean accessions. For each accession, the following multi-omics data were collected:

- Whole-genome SNP sequences: 198 × 425, 858; each variable represents SNP information.
- Rhizosphere metabolite concentrations (meta-metabolome): 198 × 263; each variable represents the concentration of a specific metabolite).
- 16S sequence data of rhizosphere microbiota genome data (microbiome): 198 × 1111; each variable represents the count of a specific microorganism.
- Soybean phenotype data: 198 × 9; biomass data such as dry weight and fresh weight, etc.

Genome data were obtained from the global soybean minicore collections (Kajiya-Kanegae et al., 2021). Microbiome and meta-metabolome data were collected from identical plots to ensure a strong relationship between the two omics datasets. Meta-metabolomic data play a key role in elucidating the complex relationships between plant phenotypes and the microbiome. Data from trials conducted in 2019 and 2020 were used in the model. Additionally, two watering treatments, control and drought, were applied to evaluate the genetic variations in response to water stress. Details of the field trials and data acquisition are provided in the Appendix.

Given the high dimensionality of omics data, particularly genomic data, linear kernels were calculated to reduce the dimensionality. The genetic relationship matrix **G** was calculated as 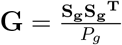, where **S**_**g**_ is the standardised genotype matrix and *P*_*g*_ is the number of SNPs. Each column of S_**g**_ is standardised such that 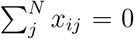 and 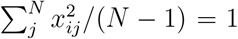 over *N* lines. The relationship matrices for the meta-metabolome and microbiome **G** were calculated similarly. Additionally, the meta-metabolome and microbiome data were scaled before being used for modeling. All statistical analyses were conducted using R version 4.4.1 (R Core Team, 2023), and figures were produced using the R package ggplot2 version 3.5.1 (Wickham H, 2016).

### Two-Step Prediction model

We propose a two-step prediction model to integrate the intermediate traits (Figure 1). The two-step prediction model is based on two hierarchical equations. Initially, the model was limited to linear regression, which solved the following multikernel linear mixed-effects model:

**Figure 1:**
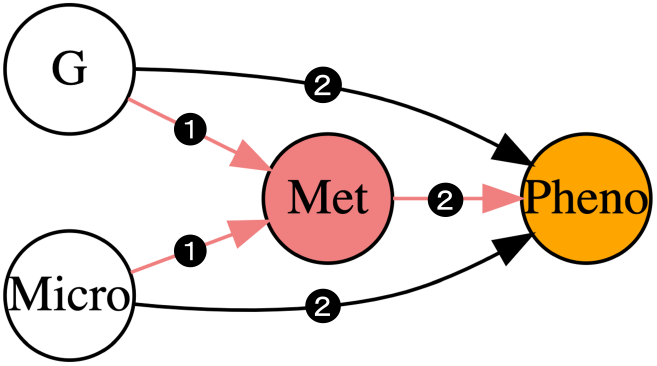
Diagram of G-Micro-Proxy model. G: Genome, Met: Meta-metabolome, Micro: Microbiome, Pheno: Phenotype. Arrows illustrate the direction of effects among omics and phenotype, demonstrating the interactions between them. Black arrows represent pathways unrelated to predicted intermediate omics traits. Red arrows represent pathways involving the intermediate traits predicted from explanatory variables (G and/or Micro). The first step predicts the meta-metabolome, and the second step predicts the phenotype. The model provides the intermediate omics traits (meta-metabolome) explained by genome and microbiome. Prediction models can be chosen as either Random Forest (RF) or Best Linear Unbiased Prediction (BLUP) for each step.

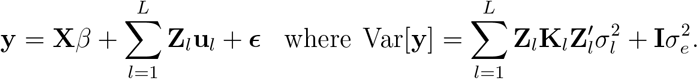

In this model, **y** represents the response variable, **X** is the fixed-effects design matrix, and *β* is the corresponding vector of fixed effects. The term **Z**_***l***_ denotes the random-effects design matrix for kernel *l*, where **u**_*l*_ represents the associated vector of the random effects. The residual error is denoted as ϵ . The variance of **y** is modeled using the covariance (kernel) matrix **K**_*l*_ for each kernel *l*, where 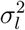 represents the variance component associated with kernel *l*. 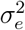 represents residual variance.

In our two-step modeling approach, we first trained a model to predict meta-metabolome profiles as intermediate traits. Subsequently, we developed a separate model to predict the phenotypes based on the predicted meta-metabolome profiles in combination with multiomics data. To evaluate the predictive abilities of the integrated models, we compared the G-Proxy and G-Micro-Proxy models. The model was based on the hypothesis that the genome and microbiome levels affect the meta-metabolome. Additionally, the meta-metabolome, genome, and microbiome levels are thought to influence the phenotype. In contrast, we assumed that the phenotype did not influence these omics levels, and that the meta-metabolome did not affect the genome or microbiome. The R package ‘RAINBOWR’ version 0.1.35 in R (Hamazaki & Iwata, 2020) was used for the computation of multi-kernel linear mixed effects model.

### G-Proxy model

First, meta-metabolome data were predicted from the genomic data and replaced with the predicted values. Secondly, the phenotype was predicted using genomic data and the predicted “proxy” metabolomic data. The remainder of this paper is structured as follows.

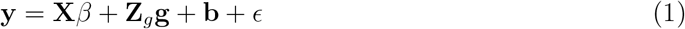

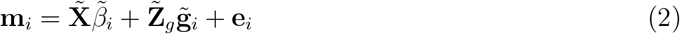

Equation (1) describes the plant phenotype prediction from the genome and intermediate omics traits. Here, **y** is the vector of phenotypes, *β* is the vector of fixed effects, and the matrix **X** is the fixed effects on the phenotypes. Vector **b** represents the regression effects of meta-metabolomic expression levels on the phenotypes. The vector **g** contains genetic effects, the matrix **Z**_**g**_ relates these genetic effects to the phenotypes, and *ϵ* is the vector of residuals.

Equation (2), for each omics feature *i* = 1, . .., *k*, describes the prediction of intermdediate omics expression levels **m**_**i**_, which compose a matrix **M**. Columns in **M** are the meta-metabolome profiles **m**_**1**_,. .., **m**_**k**_. 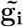 represents genetic effects, the matrix 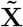 relates the fixed effects to omics expression levels, and the matrix 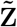 relates individuals to omics expression levels. Additionally, the vector 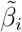 contains fixed effects, and **e**_**i**_ is the vector of the residuals.

To obtain intermediate omics values, BLUP was performed for each feature using a LeaveOne-Out (LOO) approach. The distribution assumptions for the variables were as follows:

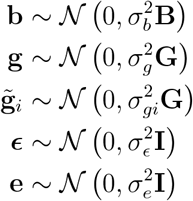

Here, **G** represents the genetic relationship matrix and **B** represents the relationship matrix derived from the predicted meta-metabolome. Matrix **B** is computed as follows:

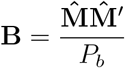

where *P*_b_ represents the total number of meta-metabolomic features. In this equation, 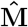 represents the meta-metabolome profiles predicted using leave-one-out cross-validation (LOOCV) based on Equation (2). Each column of 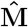 is standardised.

### G-Micro-Proxy model

We extended the model by incorporating microbiome data. This addition aims to improve the prediction accuracy and provide deeper insights into the relationships among omics layers. In the first step, the meta-metabolome is predicted using genome and microbiome data, with the resulting predicted values prepared as “proxies”. Second, the phenotype was predicted using genomic, microbial, and proxy metabolomic data. The model is as follows:

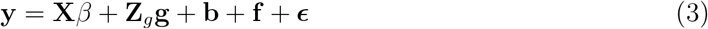

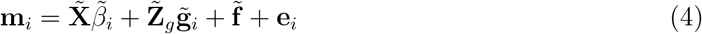

Following the G-Proxy model, Equation (3) describes the phenotype prediction from the genome, microbiome, and predicted meta-metabolome as intermediate omics traits. Equation (4) describes the meta-metabolome prediction from the genome and microbiome data.

Vectors **f** and 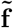 contain the regression effects of microbial expression levels. The probability distributions are defined as follows:

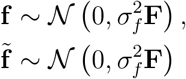

where **F** represents the microbial relationship matrix computed as 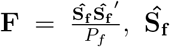 is the standardised microbial matrix, and *P*_*f*_ is the number of microbial features.

### Alternative strategies including Random Forest model

An alternative model with an ((RF) for each step. RF is a versatile machine-learning method that has been widely used in omics studies (Pfeifer et al., 2022; Qi, 2012; Chen et al., 2012). RF operates an ensemble algorithm based on randomised regression trees, where each tree is constructed from a randomly selected sample with replacement (bootstrap sample) from the training dataset, along with randomly chosen explanatory variables. The final predictions were derived by aggregating the predictions from all trees in the forest using all the explanatory variables. Its applicability is particularly notable for analyzing interconnected genomic, meta-metabolomic, and microbial datasets, providing insights into intricate relationships within biological systems. To validate the performance of the RF in this prediction task, we replaced BLUP with RF at each step of the model. The RF model can incorporate nonlinearity compared to the classic linear model, possibly contributing to higher predictive accuracy.

RF modeling was performed using the genome, microbiome, and metabolome data as inputs. Owing to the high dimensionality of genome data, which poses computational challenges for the RF model, the GRM was used as an alternative input. The efficiency of GRM in RF has recently been demonstrated (Inamori et al., 2024). Scaled values were used for microbiome and metabolome datasets, whereas for the predicted meta-metabolome, the input corresponded to 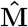.

RF was performed for each feature using leave-one-out (LOO) cross-validation for each individual, replacing the intermediate omics values with predictions. We then tested whether the predicted intermediate traits improved the accuracy of the trait predictions. Specifically, we evaluated all four model combinations (BLUP-BLUP, BLUP-RF, RF-BLUP, and RF-RF) to validate phenotype predictions. Names indicate the models used in the first and second steps. For example, RF-BLUP represents a model in which RF is used in the first step and BLUP is used in the second step. The R package “ranger” package version 0.16.0 in R (Wright & Ziegler, 2017) was used to estimate the parameters of the RF model.

### Inter-Year meta-metabolome prediction

We also demonstrated interyear predictions using the meta-metabolome. In this approach, the intermediate omics prediction model was trained using the 2020 data and tested on the 2019 data. This model generates “proxy” intermediate traits, supporting the phenotype prediction model. By leveraging this approach, the “proxy” meta-metabolome can be used to support model performance. Once the two-step prediction model was established, the need for actual intermediate omics data during prediction was eliminated, thus significantly reducing data acquisition costs.

### Simulation

In this study, we demonstrated our models using simulated datasets designed to achieve two key objectives.

First, we evaluated the effectiveness of the predicted intermediate omics data using a twostep prediction model. Specifically, we compared the performance of a model using predicted intermediate omics to that of a model using observed intermediate omics. Unlike observed omics, predicted omics traits are solely explained by omics data, and are expected to capture generalisable information that contributes to a more robust model.

Second, we investigated how a nonlinear approach improves intermediate omics predictions. To illustrate this, we generated datasets under both linear and nonlinear scenarios and applied two modeling approaches: RF and BLUP. By comparing their performances, we assessed how well the nonlinear model captured the inherent complexity of omics relationships.

Thus, we simulated omics-omics interactions under both linear and nonlinear scenarios and subsequently predicted traits using a linear model (BLUP) and a nonlinear model (RF).

We assumed that intermediate omics traits are controlled by the genome, where both the genome and the intermediate omics trait independently affect the phenotype, as described by Christensen et al. (2021). In general, a probabilistic graphical model of omics data can be illustrated using *A, B*, and *C*, as follows:

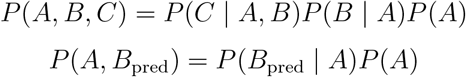

Here, we assume that *A* and *B* represent different types of omics data, and *C* represents the phenotype. The intermediate omics trait predicted by *A* was denoted as *B_pred_*.

Although this model explains certain relationships within the omics framework, it can also be extended to account for additional unknown factors, such as environmental effects, represented by *D*. The model is expressed as follows:

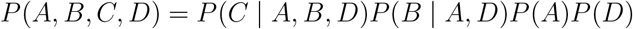

A graphical model is shown in Figure 2. The unknown factors *D* do not influence *A* or the predicted variable (*B*_*pred*_). Instead, the *D* variables directly affect *B* and *C* via independent pathways. Consequently, *B*_*pred*_ reflects the influence of *A* by isolating it from the unknown factors *D*. Assuming that *B*_*pred*_ is conditionally dependent only on *A*, it follows that *B*_*pred*_ is independent of *D*.

**Figure 2:**
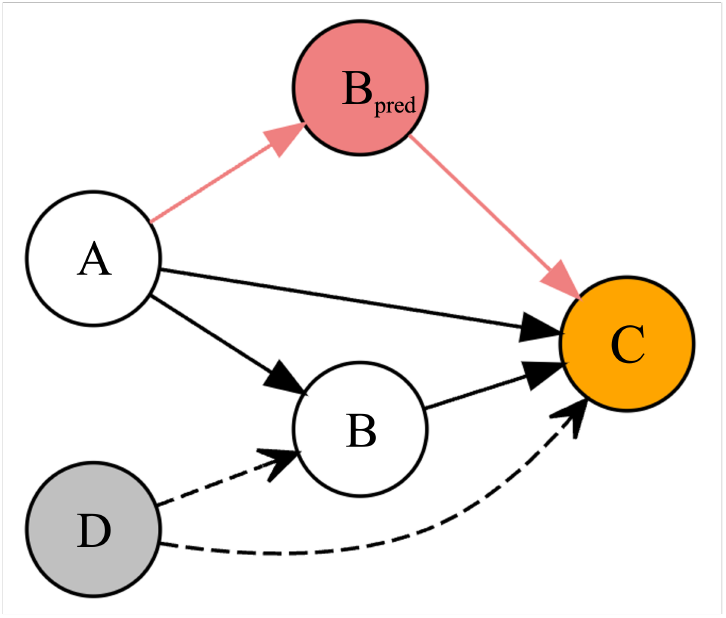
Diagram of simulations. We assume that A, B, and C represent different types of omics data, while D represents unknown factors affecting B and C. Arrows illustrate the direction of effects among omics, demonstrating the interactions between them. Black arrows represent pathways unrelated to predicted intermediate omics traits (*B*_*pred*_). Red arrows represent pathways involving *B*_*pred*_, intermediate traits predicted from explanatory variables in A. Broken arrows denote the influence of unknown factors on the traits, which are only observable in simulations. The intermediate omics traits (*B*_*pred*_) are predicted solely based on A. The model incorporates the intermediate phenotype *B*_*pred*_, which is influenced independently by the unknown factor D on B and C. *B*_*pred*_ contains only information derived from A, excluding any contribution from D

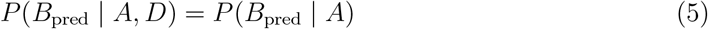

Thus, we demonstrate that building a model based on *B*_*pred*_ to remove unknown environmental factors *D* can provide a more generalised and robust modeling framework. See the Appendix for the proof of (5).

### The linear scenario

In the linear scenario, we simulated the relationships among omics data using linear combinations of matrices. Four matrices, *A, B, C*, and *D*, were generated for the interaction based on the predefined relationships. The simulation was conducted for ten iterations to generate data and facilitate predictions. Specific equations were used to model these interactions, enabling the predictive modeling of the relationships among the matrices.

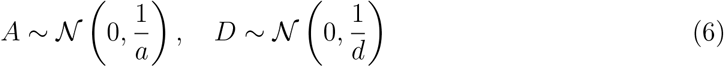

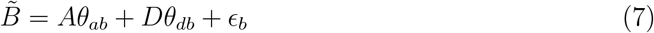

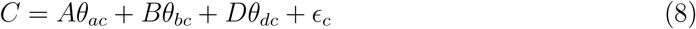

Let *A* be an *n* × *a, D* an *n* × *d*, *B* an *n* × *b, C* an *n* × *c* matrix. For computational convenience, *B* was normalised to 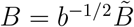 . The coefficient parameters *θ*_*ab*_,*θ*_*db*_,*θ*_*ac*_,*θ*_*bc*_,and *θ*_*dc*_ are randomly generated from a normal distribution 𝒩 (0, *I*). The noise parameters *ϵ*_*b*_ and *ϵ*_*c*_ follow 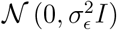. For the simulation, we set *n* = 30, *a* = 10, *b* = 10, *c* = 10, *d* = 10, and *σ*_*ϵ*_ = 0.5. The model assumes a scenario in which omics data matrices A, *B*, and *C* are used with an unknown factor *D* influencing both *B* and *C* (e.g. environmental effects on omics data and phenotypes).

### The nonlinear scenario

Next, we consider a scenario involving nonlinearities. This scenario was based on the assumption that nonlinear interactions frequently play a role in complex omics relationships. To introduce nonlinearity, we apply different functions, such as the ReLU (Rectified Linear Unit) function, Hadamard product, and sigmoid transformation, to equations (7) and (8). The equations are as follows:

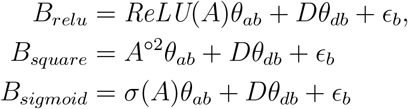

*A*^○2^ represents the Hadamard product, where each element *a*_*ij*_ of matrix *A* is squared element-wise. The term *ϵ*_*b*_ denotes the error. The ReLU transformation, denoted by *ReLU*, is defined as

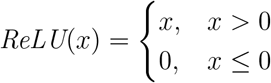

where *x* denotes an input value. The sigmoid function, denoted by *σ*(*x*), is defined as:

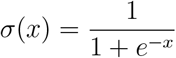

The generation of *A* and *D* follows a normal distribution as described in Equation (6). To generate the matrix *C*, the intermediate matrix *B* is selected from one of the following transformations: *B*_relu_, *B*_square_, or *B*_sigmoid_, depending on the context of the simulation.

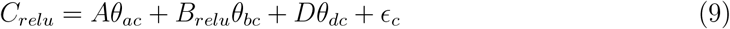

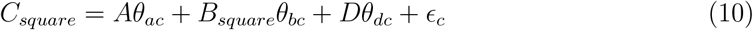

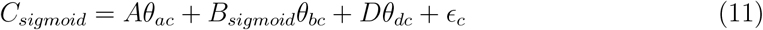

The terms *C*_relu_, *C*_square_, and *C*_sigmoid_ incorporate nonlinearity into the data generation process. The parameters were aligned with those used in linear scenarios. Furthermore, we can incorporate nonlinearities from *A* to *C* as described in the Appendix.

## 3 Results

### 3.1 Simulation Results

Simulation data were generated following the scenarios described in the previous section (linear, ReLU, square, and sigmoid) to compare the results based on nonlinearity in omics-omics relationships. For all four scenarios, we performed predictions using our proposed two-step model, in which either the BLUP or RF model was selected for each prediction task. Therefore, we compared the BLUP-BLUP, RF-BLUP, RF-RF, and BLUP-RF models. The model was validated using the leave-one-out (LOO) method for each step of the two-step prediction model. The simulations were performed for 10 iterations across all models and scenarios.

In the first step of the model, we evaluate the prediction of B from A. The data were generated under either a linear or nonlinear scenario, and predictions were made using BLUP or RF to compare prediction accuracy. The prediction results for B are shown in Figure 3. The RF model significantly outperformed the BLUP model in nonlinear scenarios (ReLU, square, and sigmoid), whereas BLUP performed better than RF in the linear scenario.

**Figure 3:**
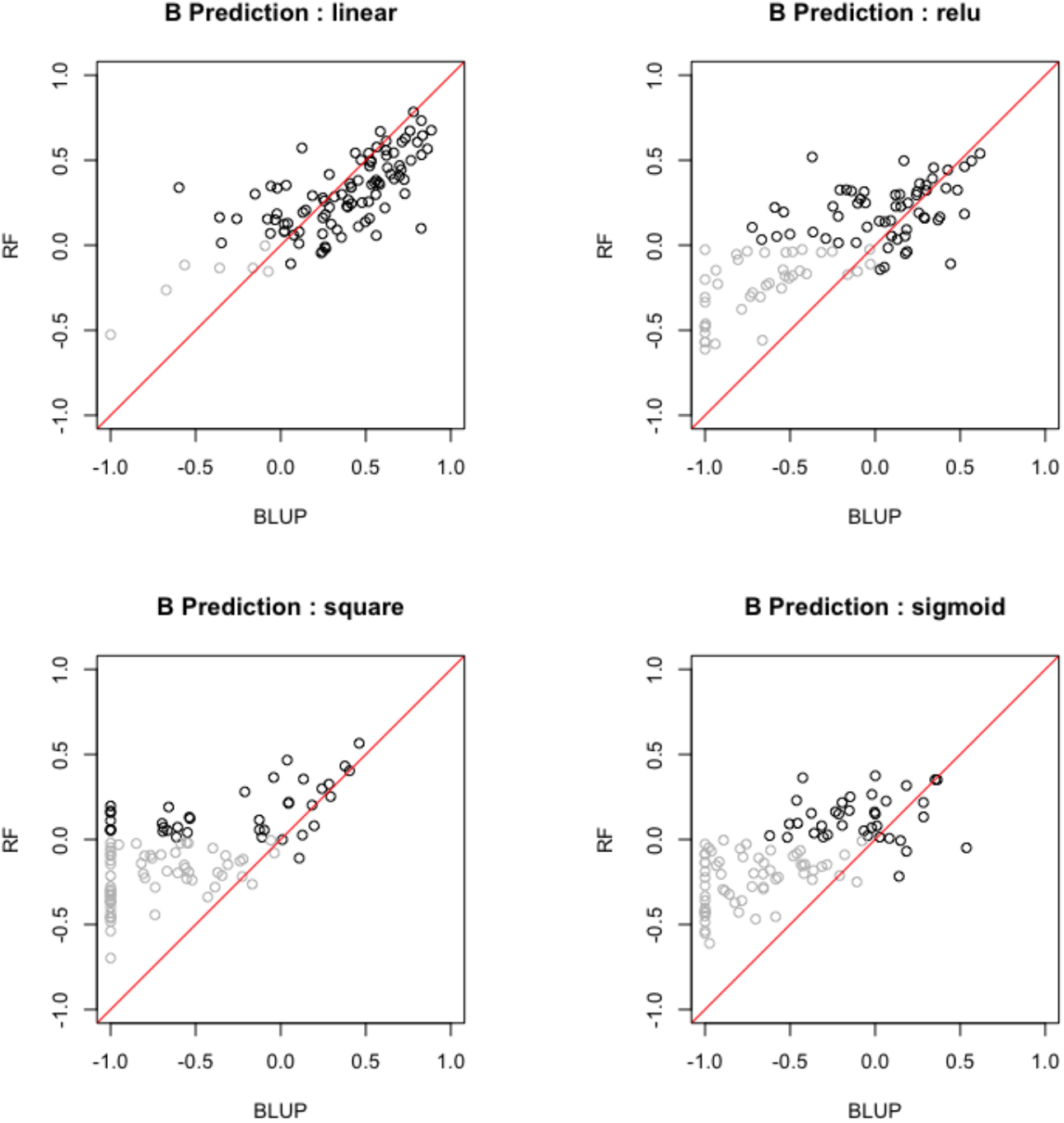
Results of the First Step: Comparison of model accuracy for B predictions using A by RF vs BLUP under different scenarios: linear model, ReLU model, square model, and sigmoid model. Each data point represents a correlation coefficient calculated via LOOCV for an individual iteration, with those where both correlation coefficients are negative shaded in gray.

In the second step, we evaluated the prediction of *C*. Specifically, we compared two models: (i) a model trained using *A* and *B*_pred_, and (ii) a model trained using *A* and the observed *B*. In both models, the prediction is applied to *A* and *B*_pred_. Figure 4 presents the BLUP-BLUP model as a representative example because it exhibits the highest accuracy. The results indicated that model (i) outperformed model (ii), suggesting that the model should be trained on the predicted values rather than the observed values when the intermediate omics trait is not available for testing.

**Figure 4:**
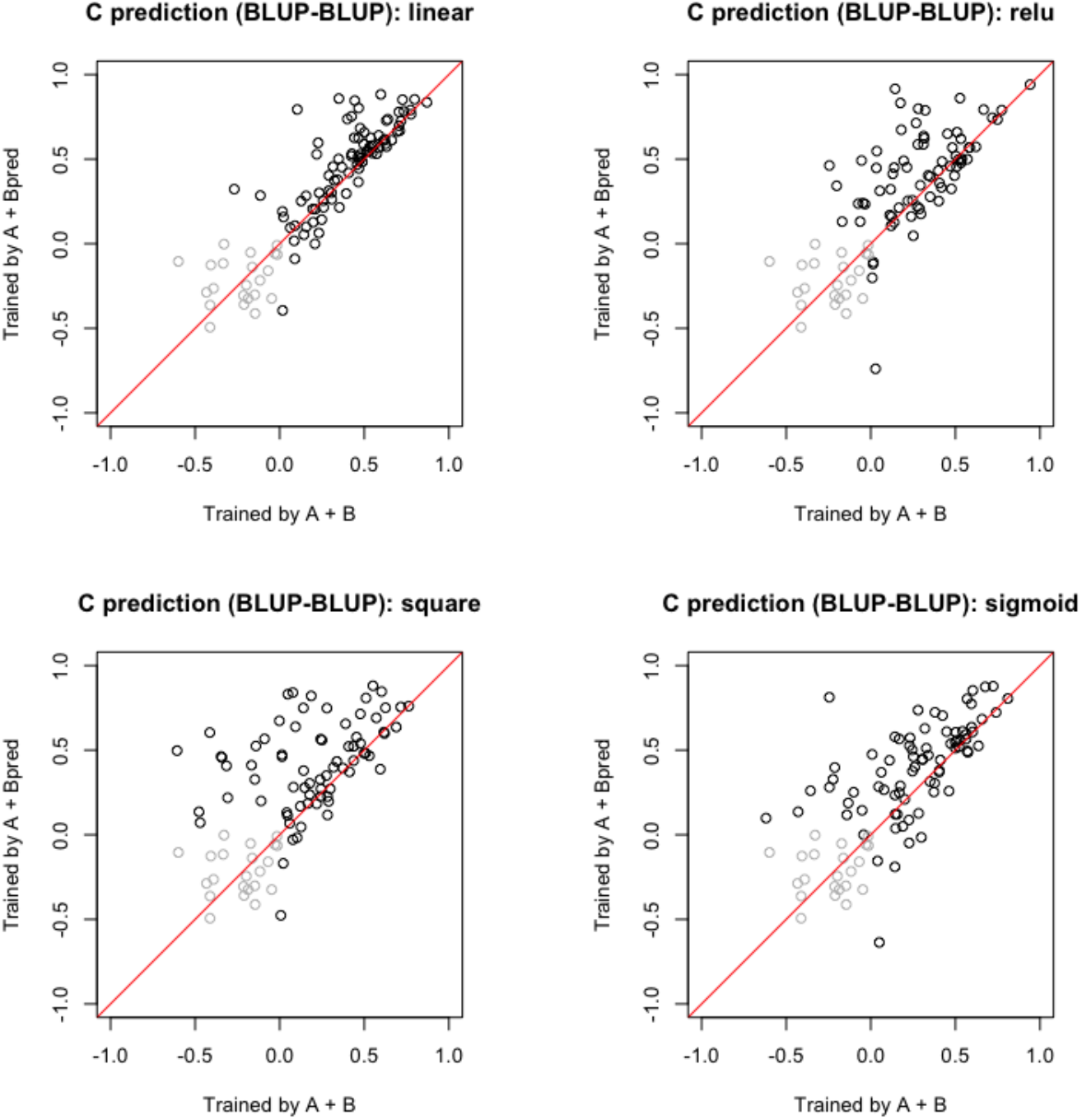
Results of the Second Step: Comparison of model accuracy for *C* predictions trained using either *A* + *B*_pred_ or *A* + *B*. Here, *B*_pred_ represents the predicted value from the first-step model using *A*. The trained models were applied to *A* + *B*_pred_ to assess prediction performance under different scenarios: linear model, ReLU model, square model, and sigmoid model. The simulations were conducted under the BLUP-BLUP model, where BLUP was used to obtain *B*_pred_ in the first step, followed by BLUP to predict *C* in the second step. Each data point represents a correlation coefficient calculated via LOOCV for an individual iteration, with those where both correlation coefficients are negative shaded in gray.

The results for all model combinations (BLUP-BLUP, RF-BLUP, RF-RF, and BLUP-RF) are shown in Figures 1S and 2S. Notably, the model generally performs better when trained with *B*_pred_ than with the observed *B* across both linear and nonlinear scenarios in both BLUP and RF models. Interestingly, in the linear and nonlinear scenarios, the BLUP-BLUP and RF-BLUP models showed comparable performances. The RF-RF model exhibited a lower accuracy but fewer outliers, demonstrating its robustness. Conversely, the BLUP-RF model had a much lower accuracy than the other models.

Simulations were further conducted for scenarios with nonlinearity between A and C. In these cases, the RF-RF model outperformed the others in nonlinear scenarios, whereas the BLUP-BLUP model performed the best in the linear scenarios. The detailed model equations are provided in the Appendix. The results are shown in Figures 3S and 4S.

### 3.2 Application Results

#### The First Step: Meta-Metabolome Prediction

In the first step of the modeling process, we predicted meta-metabolome profiles based on genome and/or microbiome data. We evaluated the RF and BLUP as predictive models. Our analysis included both intra- and inter-year models. The intra-year model was trained and tested on data from 2019, whereas the inter-year model was trained on 2020 data and tested on 2019 data. To validate the models, we used LOOCV under both the control and drought conditions. Genome and/or microbiome data were used as model inputs to assess the predictive power of the meta-metabolome profiles. We compared three scenarios: G (genome only), micro (microbiome only), and G+Micro (a combination of genome and microbiome data).

The results of the intra-year predictions, presented in Figure 5, indicate that incorporating genome and microbiome data improves the prediction accuracy. Notably, the microbiome data contributed significantly to the predictive performance, with this effect being more pronounced under drought conditions. Additionally, according to the prediction patterns, the G+Micro model aligned more strongly with the micromodel than with the G model (Figure 5S).

**Figure 5:**
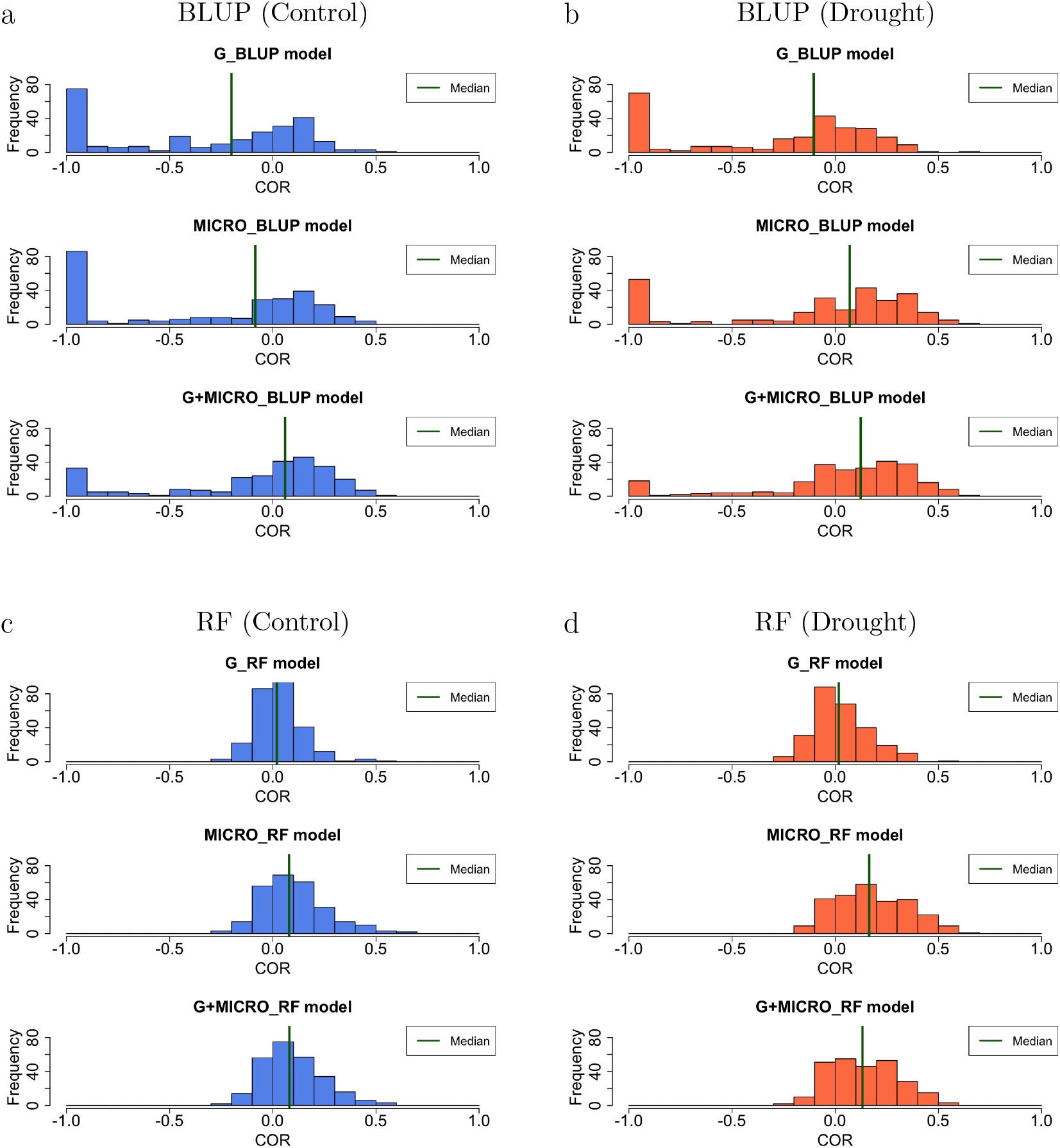
The accuracy of each meta-metabolome prediction (correlation coefficients) is shown in histogram. The model was trained and tested using 2019 data. a) BLUP-Control b) BLUP-Drought c) RF-Control d) RF-Drought. For each, the input of the model is Genome (Top), Microbiome (Middle), both Genome and Microbiome (Bottom). The green vertical line represents the median.

The results of the interyear predictions are presented in Figures 6S and 7S. Consistent with the intra-year model, incorporating genome and microbiome data improved the prediction accuracy. Detailed results are presented in Table 1S.

**Figure 6:**
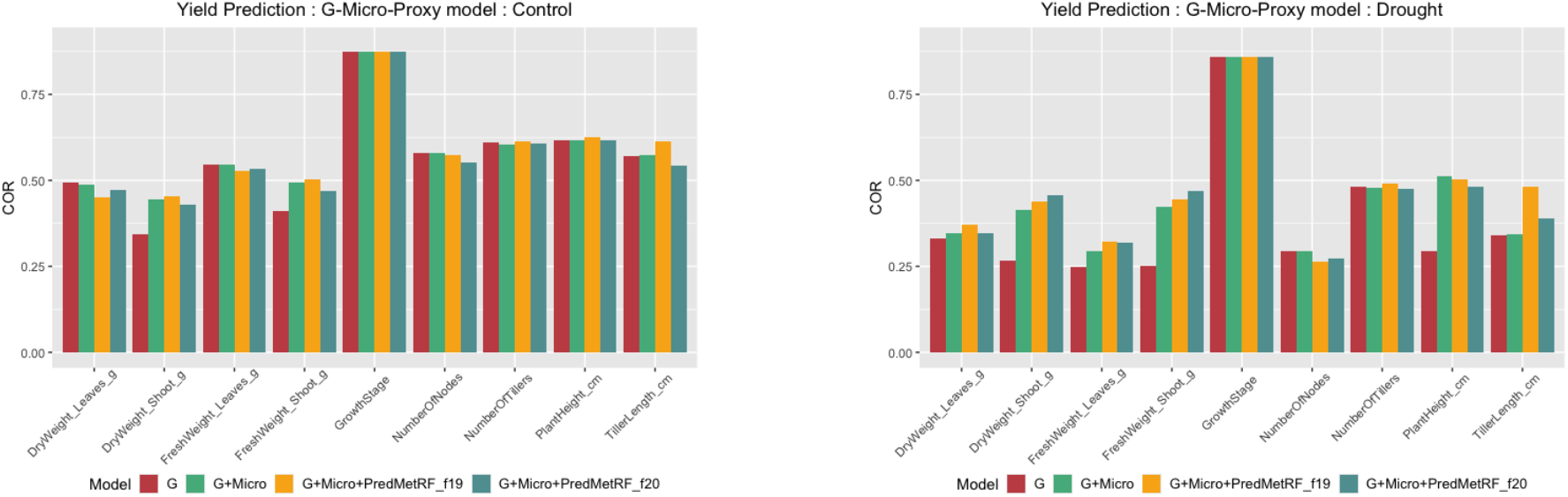
Phenotype predictions under control and drought conditions. The accuracies of each phenotype prediction (correlation coefficients) are shown. The G-Micro-Proxy models were used for phenotype predictions. G stands for Genome, Met for Meta-metabolome, and Micro for Microbiome. The first step consisted of predicting intermediate meta-metabolomic characteristics using RF, followed by BLUP for the prediction of the next phenotype. The legend illustrates the model input. When the input is G (red), the model follows a classic GBLUP approach. G + Micro (green) represents a multi-kernel BLUP incorporating both genome and microbiome data. The meta-metabolome was predicted either within the same year (19: Intra-year, yellow) or across years (20: Inter-year, blue). Each model was evaluated using leave-one-out cross-validation (LOOCV), and correlation coefficients were calculated accordingly.

In addition, we compared BLUP and RF for meta-metabolome predictions. In both the intra-and interyear predictions, as shown in Figure 5 and 6S, the RF model outperformed BLUP. In particular, in the intra-year model, BLUP predictions exhibited frequent outliers, whereas RF demonstrated a more stable prediction pattern. Inter-year, the prediction patterns of the RF model indicated that the G + Micro model was more aligned with the Micro model than the G model, whereas BLUP indicated that the G model was closer to the G+ Micro model (Figure 7S).

**Figure 7:**
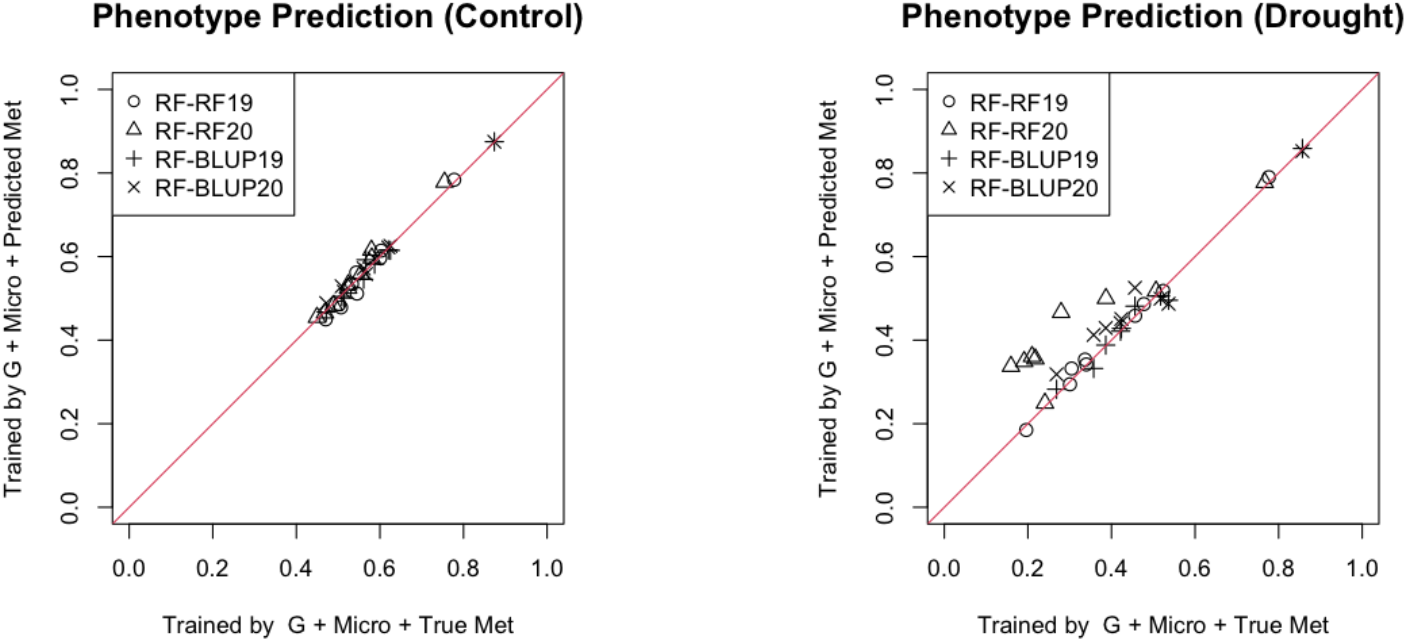
Comparison of the correlation coefficients for phenotype predictions made using predicted meta-metabolome data versus true (observed) meta-metabolome data. The models were trained on either predicted or observed meta-metabolome data and then tested on predicted meta-metabolome data. In both cases, Genome and Microbiome were given for the inputs by default. The legend shows Model names following the format ‘F-SYY,’ where F, S represent the methods (BLUP or RF), and YY indicates the training period (19 for intra-year or 20 for inter-year). Each data point represents a different phenotypic trait.

#### The Second Step: Phenotype Prediction

For phenotype prediction, either BLUP or RF were used. Given RF’s superior performance of RF in the first step of the simulation and application, leveraging RF-predicted meta-metabolome data was deemed a robust approach. Consequently, RF was used in the first step to predict intermediate meta-metabolomic traits, followed by BLUP in the second step for the final phenotype prediction, allowing direct comparison with conventional GBLUP. Additionally, four prediction patterns—BLUP-BLUP, BLUP-RF, RF-BLUP, and RF-RF—were evaluated (Figure 9S). All models were validated using leave-one-out cross-validation (LOOCV), and their predictive accuracies were assessed based on correlation coefficients.

We compared the G-Proxy and G-Micro-Proxy models to evaluate their predictive performances. The results for the G-Micro-Proxy model are presented in Figure 6, and those for the G-Proxy model are shown in Figure 12S. Our analysis demonstrates that integrating microbiome data within a two-step framework significantly improves predictive accuracy, particularly under drought conditions. In the G-Micro-Proxy model, the incorporation of RF-predicted proxy data into BLUP enhanced phenotype estimation. This improvement was observed in both the intra- and inter-year periods, as shown in Figure 6. Notably, the prediction accuracy of shoot and leaf weights was significantly enhanced. Conversely, under the control conditions, both the G-Proxy and G-Micro-Proxy models exhibited predictive performances comparable to those of GBLUP.

For inter-year modeling, the phenotype predictions for 2019 were improved by utilising proxy meta-metabolome data predicted using a model trained on 2020 data (Figure 6). This finding suggests that once the two-step prediction model is established, phenotype predictions can be performed without the need for new meta-metabolome data collection. The detailed results of the G-Proxy and G-Micro-Proxy models are presented in Tables 2S and 3S, respectively. A comparison of all models across traits is shown in Figures 10S and 11S.

Futhermore,, phenotype predictions using proxy (predicted) versus true (observed) meta-metabolome data were examined. The models were trained on either the predicted or observed meta-metabolome data and subsequently tested on the predicted meta-metabolome data. In both cases, in addition to the meta-metabolome, genome, and microbiome data were used as inputs (Figure 7). The results indicate that the models trained on the predicted meta-metabolome data outperformed those trained on the true meta-metabolome data, particularly in interyear settings.

## 4 Discussion

### Nonlinearity in Intermediate Omics Traits

The simulation results show that the RF model is highly effective for omics-omics modeling with nonlinearity (Figure 3). Consistent with this result, the application analysis demonstrated that the RF model outperformed BLUP in meta-metabolome prediction (Figure 5 and 6S). Additionally, the prediction patterns indicated that the microbiome was a mejor factor for meta-metabolome prediction, particularly in the RF model (Figure 6S and 7S). This suggests that nonlinearity exists in the relationships within omics. Some microbiome and metabolome data exhibited discrete patterns (e.g. presence/absence), which also implies that the nonlinear RF model was effective in capturing these patterns. These results demonstrate that the accuracy of phenotype prediction can be improved by incorporating predicted metabolites, and that the model can be improved by introducing nonlinearity through RF.

### Advantages of Two-Step Modeling Approach

Through simulation and application, the two-step modeling approach, which integrates meta-metabolome data as an intermediate layer, has been proven to be effective for predicting phenotypes. In particular, under drought conditions, RF was advantageous for modeling the relationships between omics in the first step, and applying BLUP in the second step improved phenotype prediction (Figure 8S and 9S). In contrast, under control conditions, BLUP performed well in both the first and second steps, suggesting that the microbiome contributed less to the trait, leading to relatively simpler interactions among omics with an accuracy comparable to that of conventional GBLUP. These results imply that in the presence of complex environmental factors, RF may better capture omics relationships through nonlinear modeling. These interactions could be further elucidated through simulations and research across diverse scenarios.

### Microbiome Contribution to Phenotype Prediction

Under drought conditions, the metabolites predicted by RF were used as intermediate phenotypes, achieving better accuracy than GBLUP in predicting the shoots and leaves. In contrast, under control conditions, the accuracy of phenotype prediction was comparable to that of GBLUP (Figures 6 and 12S). Our results indicate a strong association between the meta-metabolome and microbiome under dry conditions. Under drought stress, the G-Micro-Proxy model significantly outperformed the traditional GBLUP model, confirming that the important role of the microbiome in regulating plant phenotypic traits (Figure 6). This finding highlights the importance of adopting a two-step model that incorporates microbiome information into a phenotype prediction framework. This improved model could facilitate the development of breeding strategies under various environmental conditions.

### Isolating Genetic and Microbial Effects in Meta-metabolome data

This study emphasises the importance of isolating and analyzing intermediate omics traits from confounding environmental factors. Through simulations, we showed that data between omics may contain unknown environmental effects that complicate predictive work. In simulation and application, we have demonstrated that the two-step strategy can eliminate these undesirable effects among omics data to extract “proxy” intermediate omics traits. Importantly, the “proxy” omics traits can be used to improve predictive accuracy (Figure 4 and 7). This approach also suggests the possibility of investigating genetic and microbial determinants by isolating the relevant genetic signals from confounding environmental effects. This insight not only deepens our knowledge of genetics, but also allows us to better understand the functional role of microbial communities in metabolic processes and overall plant health, thus justifying further investigation. However, further biological validation is still required in this regard.

### Inter-Year Predictive Consistency

The success of the inter-year modeling demonstrates that our model can effectively exclude environmental factors from the model and extract only genetically interesting information as “proxy” omics traits. Furthermore, this suggests that once the prediction framework is established, phenotypes can be predicted without collecting new meta-metabolome data. This approach effectively reduces the cost of obtaining meta-metabolome data, which is especially costly, and provides a practical solution for omics-combined studies (Figure 6).

## 5 Conclusion

This study advances the predictive modeling of intermediate omics traits by integrating genomic, microbiome, and meta-metabolomic data. A comparison between the Random Forest (RF) and Best Linear Unbiased Prediction (BLUP) underscores the limitations of traditional linear models in capturing nonlinear relationships, particularly under drought conditions. A key innovation of this work is the two-step prediction framework, which utilises a “proxy” meta-metabolome to reduce data acquisition costs. Incorporating RF into this framework introduces a novel methodological approach, whereas using intermediate omics as a “proxy” provides deeper insights into biological mechanisms. By isolating intermediate omics traits from environmental variability, this framework enhances our genetic understanding of these traits and provides targeted breeding strategies. Future research should explore factor analysis to further refine the latent structure of multi-omics data, thereby improving both prediction accuracy and biological interpretation.

## Data availability

Data supporting the findings of this study are available from the corresponding author, H.I., upon reasonable request. All source codes are available from the repository in GitHub: https://github.com/Yoska393/Twostep. The metabolome data are available from the RIKEN DropMet website (http://prime.psc.riken.jp/menta.cgi/prime/drop_index; ID: DM0071, DM0072).

## Acknowledgement

We thank Yutaka Yamada (RIKEN CSRS) for informatics support. We are grateful to the technical staff at the Arid Land Research Center, Tottori University, and Izumi Higashida.

