## Appendix for "Integration of Proxy Intermediate Omics traits into a Nonlinear Two-Step model for accurate phenotypic prediction"

[1] *Graduate School of Agricultural and Life Sciences, Univ. of Tokyo, Tokyo, Japan* [2] *Université Paris-Saclay, AgroParisTech, INRAE, UMR MIA Paris-Saclay, 91120, Palaiseau, France* [3] *Université Paris-Saclay, INRAE, CNRS, AgroParisTech, GQE - Le Moulon, 91190, Gif-sur-Yvette, France.* [4] *Institute for Agro-Environmental Sciences, The National Agriculture and Food Research Organization, Tsukuba, Ibaraki,Japan* [5] *Arid Land Research Center, Tottori University, Tottori, Japan* [6] *Graduate School of Bioagricultural Sciences, Nagoya Univ., Nagoya, Japan* [7] *Faculty of Life and Environmental Sciences, Tsukuba Plant Innovation Research Center, Univ. of Tsukuba, Tsukuba, Ibaraki, Japan* [8] *Faculty of Food Nutritional Sciences, Toyo University, Asaka, Saitama, Japan* [9] *Soybean and Field Crop Applied Genomics Research Unit, Institute of Crop Science, National Agriculture and Food Research Organization, Tsukuba, Ibaraki, Japan* [10] *RIKEN Center for Sustainable Resource Science, RIKEN, Tsurumi-ku, Yokohama, Japan* [11] *RIKEN BioResource Research Center, RIKEN, Tsukuba, Ibaraki, Japan* [12] *RIKEN Center for Sustainable Resource Science, RIKEN, Tsukuba, Ibaraki, Japan*

### 1.1 Supplementary Figures and Tables

Table 1S: Meta-metabolome prediction. Summary of correlation coefficient in fitting for all metabolites

| | Control | | | | | | Drought | | | | | |
|---|---|---|---|---|---|---|---|---|---|---|---|---|
| | BLUP | | | RF | | | BLUP | | | RF | | |
| | G | Micro | G+Micro | G | Micro | G+Micro | G | Micro | G+Micro | G | Micro | G+Micro |
| | Intra-Year Met Prediction | | | | | | | | | | | |
| Min. | -1.000 | -1.000 | -1.000 | -0.299 | -0.196 | -0.252 | -1.000 | -1.000 | -1.000 | -0.291 | -0.263 | -0.217 |
| 1st Qu. | -0.991 | -0.347 | -0.076 | -0.055 | 0.028 | 0.002 | -0.961 | -1.000 | -0.216 | -0.039 | -0.016 | -0.006 |
| Median | -0.104 | 0.070 | 0.123 | 0.018 | 0.165 | 0.133 | -0.200 | -0.085 | 0.059 | 0.020 | 0.078 | 0.080 |
| Mean | -0.311 | -0.116 | 0.033 | 0.032 | 0.178 | 0.139 | -0.353 | -0.328 | -0.098 | 0.030 | 0.102 | 0.098 |
| 3rd Qu. | 0.069 | 0.269 | 0.287 | 0.107 | 0.313 | 0.252 | 0.076 | 0.125 | 0.198 | 0.086 | 0.182 | 0.179 |
| Max. | 0.601 | 0.600 | 0.600 | 0.540 | 0.605 | 0.593 | 0.522 | 0.478 | 0.578 | 0.509 | 0.612 | 0.594 |
| | Inter-Year Met Prediction | | | | | | | | | | | |
| Min. | -0.458 | -0.415 | -0.234 | -0.210 | -0.217 | -0.267 | -0.509 | -0.645 | -0.412 | -0.153 | -0.286 | -0.158 |
| 1st Qu. | -0.031 | -0.074 | -0.031 | -0.033 | -0.043 | -0.018 | -0.005 | -0.128 | -0.000 | -0.020 | -0.036 | 0.004 |
| Median | 0.020 | -0.012 | 0.021 | 0.018 | 0.030 | 0.052 | 0.044 | -0.035 | 0.047 | 0.046 | 0.038 | 0.080 |
| Mean | 0.038 | -0.025 | 0.040 | 0.042 | 0.040 | 0.062 | 0.061 | -0.062 | 0.061 | 0.061 | 0.054 | 0.099 |
| 3rd Qu. | 0.102 | 0.037 | 0.104 | 0.111 | 0.111 | 0.135 | 0.5 | 0.021 | 0.5 | 0.119 | 0.122 | 0.160 |
| Max. | 0.494 | 0.183 | 0.396 | 0.487 | 0.545 | 0.485 | 0.534 | 0.134 | 0.534 | 0.522 | 0.654 | 0.636 |

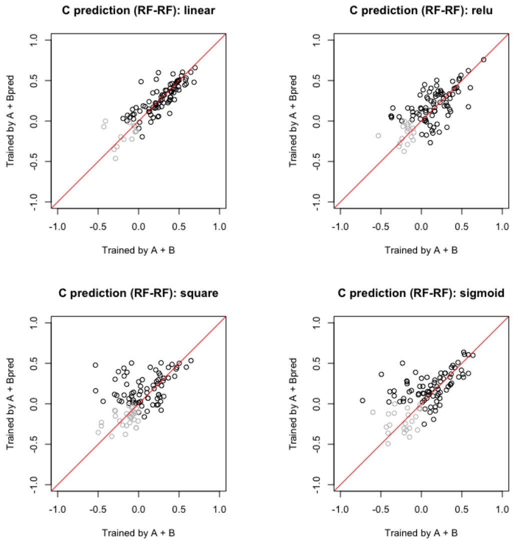


(a) RF-RF model

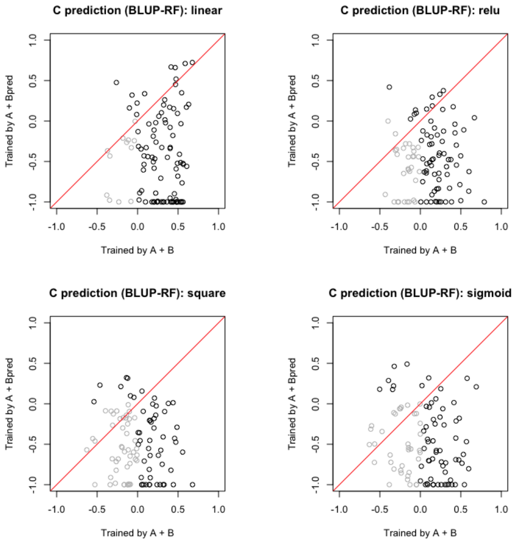


(b) BLUP-RF model

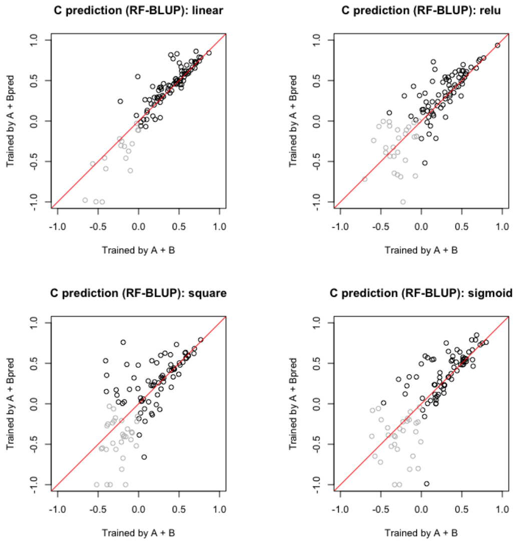


(c) RF-BLUP model

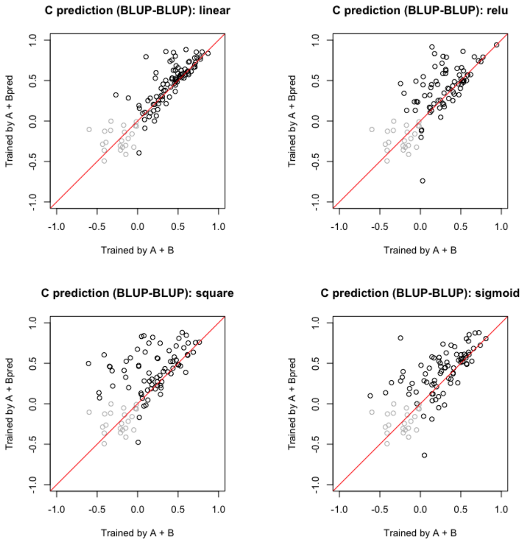


(d) BLUP-BLUP model

Figure 1S: Simulation Step 2: Comparison of model accuracy for $C$ predictions trained using either $A+B_{\text{pred}}$ or $A+B$. Here, $B_{\text{pred}}$ represents the predicted value from the first-step model using $A$. The model incorporates nonlinear interactions from $A$ to $B$. The trained models were applied to $A+B_{\text{pred}}$ to assess prediction performance under different scenarios: linear model, ReLU model, square model, and sigmoid model. Each point represents a correlation coefficient calculated via LOOCV for an individual iteration. Each data point represents a correlation coefficient calculated via LOOCV for an individual iteration, with those where both correlation coefficients are negative shaded in gray.

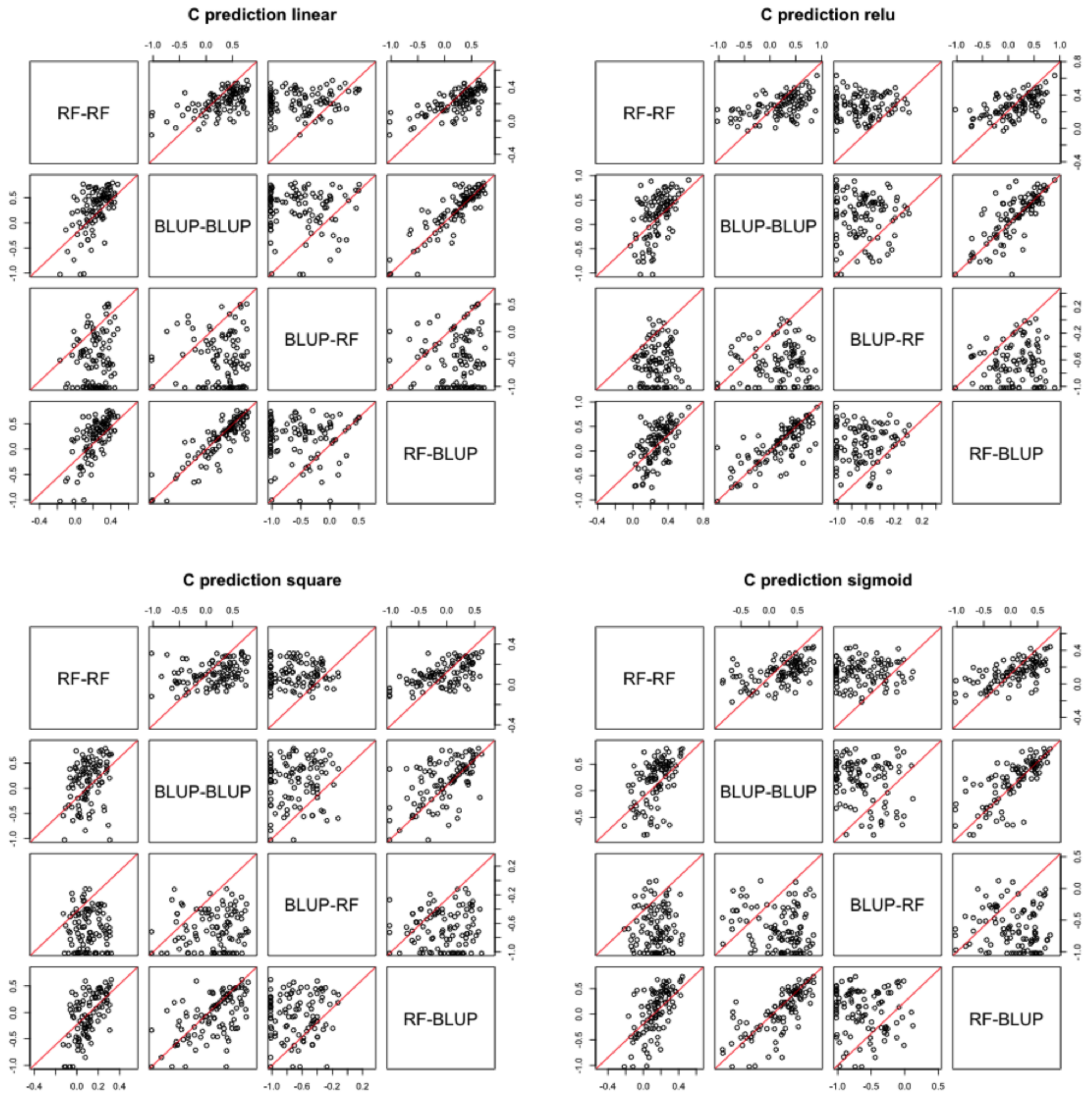


Figure 2S: Simulation Step 2: Model Comparison (BLUP-BLUP, BLUP-RF, RF-BLUP, and RF-RF) for $C$ predictions trained using $A + B_{\text{pred}}$. The model incorporates nonlinear interactions from $A$ to $B$. The models were compared under different scenarios: linear, ReLU, square, and sigmoid models. The correlation coefficient was calculated via LOOCV for each individual iteration.

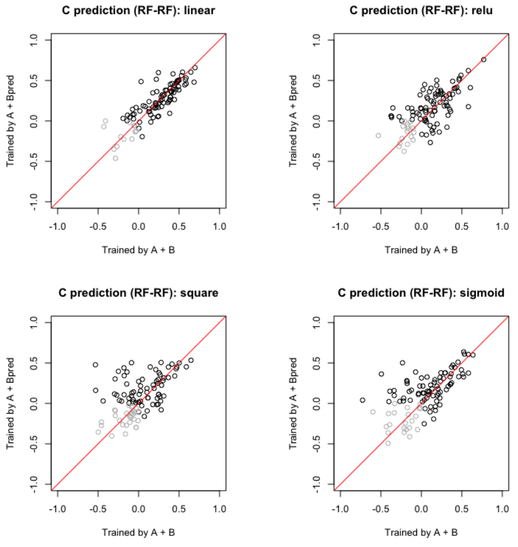


(a) RF-RF model

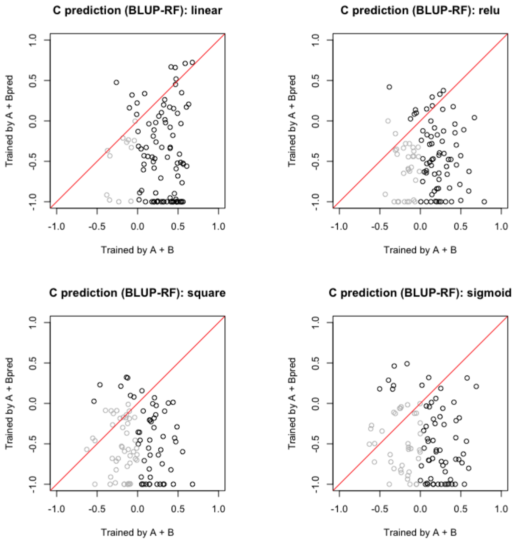


(b) BLUP-RF model

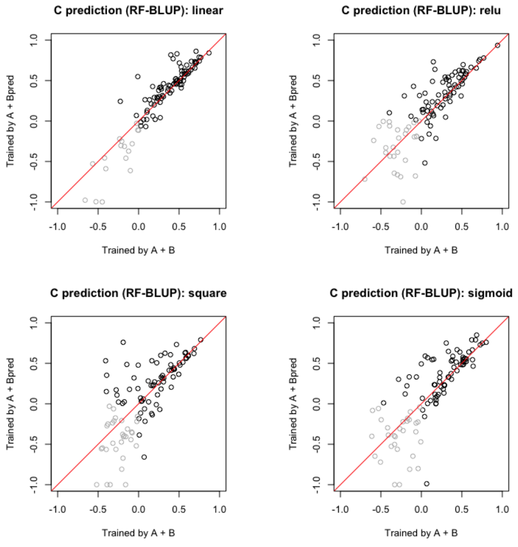


(c) RF-BLUP model

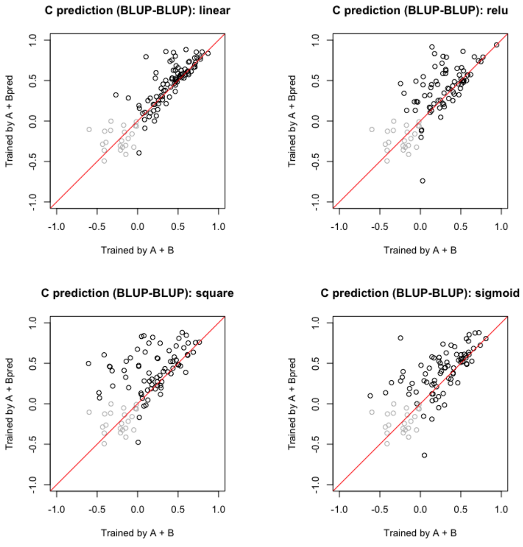


(d) BLUP-BLUP model

Figure 3S: Simulation Step 2: Comparison of model accuracy for $C$ predictions trained using either $A + B_{\text{pred}}$ or $A + B$. Here, $B_{\text{pred}}$ represents the predicted value from the first-step model using $A$. The model incorporates nonlinear interactions from $A$ to $C$, as well as from $A$ to $B$. The trained models were applied to $A + B_{\text{pred}}$ to assess prediction performance under different scenarios: linear model, ReLU model, square model, and sigmoid model. Each point represents a correlation coefficient calculated via LOOCV for an individual iteration.

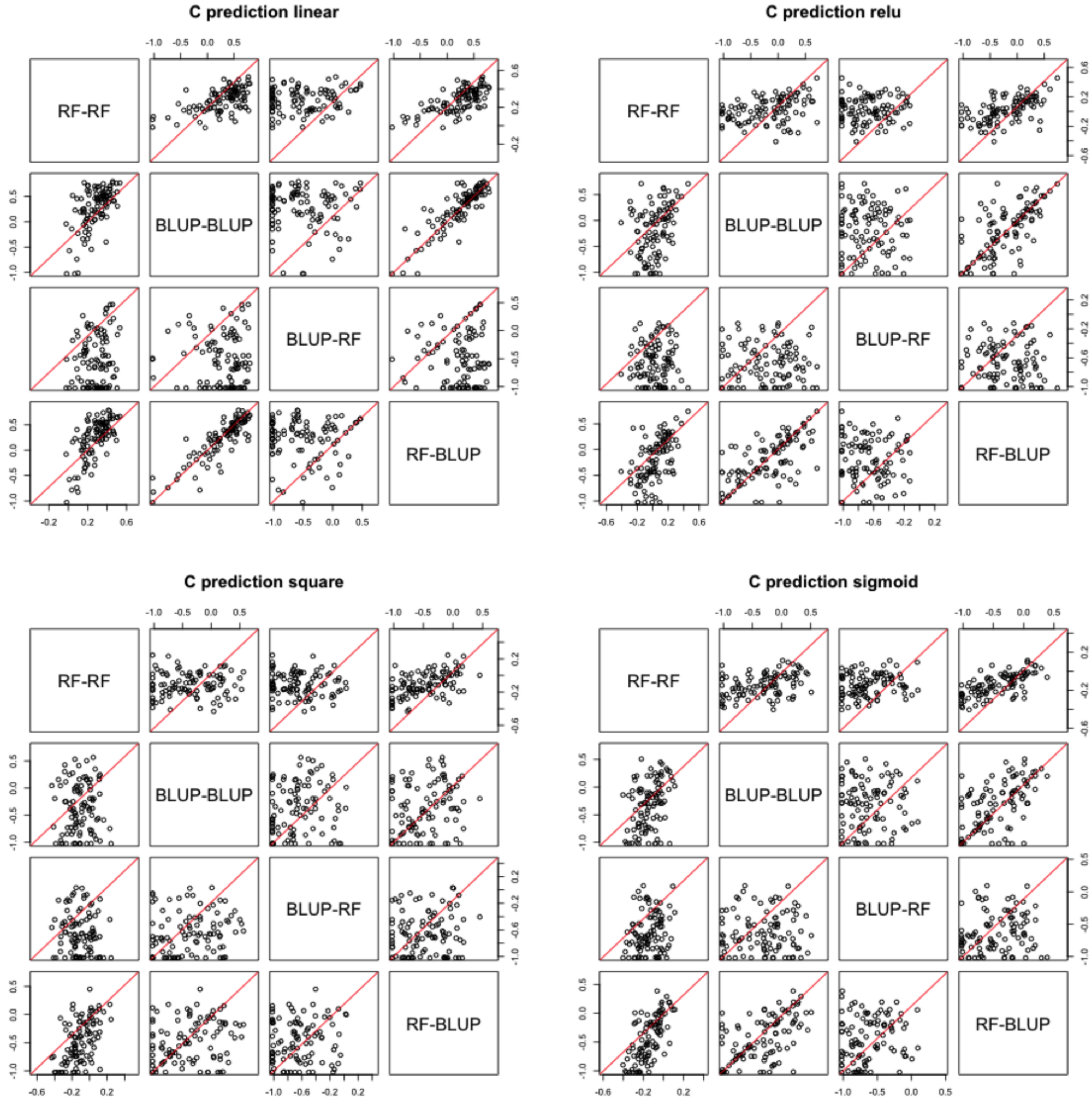


Figure 4S: Simulation Step 2: Model Comparison (BLUP-BLUP, BLUP-RF, RF-BLUP, and RF-RF) for $C$ predictions trained on $A+B_{\text{pred}}$. The model incorporates nonlinear interactions from $A$ to $C$, as well as from $A$ to $B$. The models were evaluated under different scenarios: linear, ReLU, square, and sigmoid. The correlation coefficient was calculated using LOOCV for each individual iteration.

Table 2S: All results in G-Proxy model

| Trait | Condition | Step-2 | G | G-ProxyPred | | | |
|---|---|---|---|---|---|---|---|
| | | | | **B19** | **B20** | **RF19** | **RF20** |
| DryWeight_Leaves_g | Drought | BLUP | 0.330 | 0.313 | 0.332 | 0.377 | 0.358 |
| | | RF | 0.324 | 0.380 | 0.354 | 0.424 | 0.319 |
| | Control | BLUP | 0.494 | 0.546 | 0.514 | 0.470 | 0.504 |
| | | RF | 0.492 | 0.571 | 0.542 | 0.481 | 0.498 |
| DryWeight_Shoot_g | Drought | BLUP | 0.266 | 0.253 | 0.266 | 0.261 | 0.266 |
| | | RF | 0.243 | 0.283 | 0.254 | 0.250 | 0.185 |
| | Control | BLUP | 0.342 | 0.456 | 0.426 | 0.392 | 0.389 |
| | | RF | 0.399 | 0.470 | 0.457 | 0.378 | 0.408 |
| FreshWeight_Shoot_g | Drought | BLUP | 0.253 | 0.241 | 0.253 | 0.249 | 0.253 |
| | | RF | 0.207 | 0.236 | 0.225 | 0.228 | 0.163 |
| | Control | BLUP | 0.411 | 0.510 | 0.478 | 0.443 | 0.444 |
| | | RF | 0.448 | 0.521 | 0.512 | 0.434 | 0.473 |
| FreshWeight_Leaves_g | Drought | BLUP | 0.248 | 0.236 | 0.251 | 0.336 | 0.290 |
| | | RF | 0.259 | 0.326 | 0.268 | 0.388 | 0.294 |
| | Control | BLUP | 0.547 | 0.571 | 0.570 | 0.519 | 0.561 |
| | | RF | 0.551 | 0.589 | 0.590 | 0.529 | 0.558 |
| GrowthStage | Drought | BLUP | 0.860 | 0.843 | 0.852 | 0.854 | 0.857 |
| | | RF | 0.810 | 0.799 | 0.788 | 0.800 | 0.790 |
| | Control | BLUP | 0.875 | 0.875 | 0.875 | 0.875 | 0.875 |
| | | RF | 0.816 | 0.827 | 0.823 | 0.821 | 0.813 |
| NumberOfNodes | Drought | BLUP | 0.295 | 0.290 | 0.285 | 0.288 | 0.298 |
| | | RF | 0.197 | 0.217 | 0.242 | 0.221 | 0.196 |
| | Control | BLUP | 0.580 | 0.596 | 0.565 | 0.606 | 0.593 |
| | | RF | 0.569 | 0.614 | 0.584 | 0.566 | 0.545 |
| NumberOfTillers | Drought | BLUP | 0.480 | 0.490 | 0.551 | 0.507 | 0.505 |
| | | RF | 0.498 | 0.548 | 0.531 | 0.513 | 0.524 |
| | Control | BLUP | 0.610 | 0.619 | 0.610 | 0.640 | 0.617 |
| | | RF | 0.628 | 0.648 | 0.625 | 0.635 | 0.621 |
| PlantHeight_cm | Drought | BLUP | 0.293 | 0.340 | 0.243 | 0.323 | 0.288 |
| | | RF | 0.247 | 0.260 | 0.213 | 0.238 | 0.245 |
| | Control | BLUP | 0.615 | 0.633 | 0.615 | 0.619 | 0.618 |
| | | RF | 0.627 | 0.668 | 0.619 | 0.622 | 0.617 |
| TillerLength_cm | Drought | BLUP | 0.340 | 0.419 | 0.342 | 0.480 | 0.404 |
| | | RF | 0.450 | 0.506 | 0.438 | 0.502 | 0.464 |
| | Control | BLUP | 0.570 | 0.572 | 0.569 | 0.596 | 0.541 |
| | | RF | 0.577 | 0.596 | 0.577 | 0.583 | 0.564 |
| **Sum (Mean)** | Drought | BLUP | 0.374 | 0.381 | 0.375 | 0.408 | 0.391 |
| | | RF | 0.359 | 0.395 | 0.368 | 0.396 | 0.353 |
| | Control | BLUP | 0.560 | 0.598 | 0.580 | 0.573 | 0.571 |
| | | RF | 0.567 | 0.612 | 0.592 | 0.561 | 0.566 |

Table 3S: All results in G + Micro +Met model

| Trait | Condition | Step-2 | G+Micro | G-Micro-ProxyPred | | | |
|---|---|---|---|---|---|---|---|
| | | | | B19 | B20 | RF19 | RF20 |
| DryWeight_Leaves_g | Drought | BLUP | 0.346 | 0.326 | 0.339 | 0.371 | 0.347 |
| | | RF | 0.369 | 0.326 | 0.371 | 0.330 | 0.368 |
| | Control | BLUP | 0.487 | 0.542 | 0.505 | 0.451 | 0.472 |
| | | RF | 0.507 | 0.547 | 0.534 | 0.473 | 0.488 |
| DryWeight_Shoot_g | Drought | BLUP | 0.413 | 0.400 | 0.413 | 0.440 | 0.457 |
| | | RF | 0.350 | 0.333 | 0.305 | 0.354 | 0.365 |
| | Control | BLUP | 0.446 | 0.522 | 0.464 | 0.455 | 0.431 |
| | | RF | 0.446 | 0.472 | 0.486 | 0.456 | 0.463 |
| FreshWeight_Shoot_g | Drought | BLUP | 0.424 | 0.405 | 0.424 | 0.446 | 0.469 |
| | | RF | 0.318 | 0.328 | 0.303 | 0.338 | 0.353 |
| | Control | BLUP | 0.493 | 0.563 | 0.507 | 0.502 | 0.469 |
| | | RF | 0.492 | 0.506 | 0.540 | 0.477 | 0.487 |
| FreshWeight_Leaves_g | Drought | BLUP | 0.293 | 0.255 | 0.287 | 0.322 | 0.320 |
| | | RF | 0.293 | 0.243 | 0.286 | 0.279 | 0.348 |
| | Control | BLUP | 0.545 | 0.568 | 0.560 | 0.528 | 0.533 |
| | | RF | 0.550 | 0.573 | 0.574 | 0.510 | 0.522 |
| GrowthStage | Drought | BLUP | 0.860 | 0.856 | 0.853 | 0.859 | 0.858 |
| | | RF | 0.794 | 0.796 | 0.785 | 0.786 | 0.777 |
| | Control | BLUP | 0.875 | 0.875 | 0.875 | 0.875 | 0.874 |
| | | RF | 0.786 | 0.811 | 0.817 | 0.784 | 0.779 |
| NumberOfNodes | Drought | BLUP | 0.293 | 0.273 | 0.288 | 0.263 | 0.272 |
| | | RF | 0.235 | 0.238 | 0.222 | 0.190 | 0.254 |
| | Control | BLUP | 0.580 | 0.565 | 0.596 | 0.575 | 0.552 |
| | | RF | 0.540 | 0.575 | 0.581 | 0.531 | 0.534 |
| NumberOfTillers | Drought | BLUP | 0.477 | 0.476 | 0.552 | 0.490 | 0.475 |
| | | RF | 0.506 | 0.520 | 0.531 | 0.510 | 0.510 |
| | Control | BLUP | 0.604 | 0.602 | 0.578 | 0.612 | 0.609 |
| | | RF | 0.608 | 0.621 | 0.620 | 0.607 | 0.619 |
| PlantHeight_cm | Drought | BLUP | 0.512 | 0.443 | 0.458 | 0.504 | 0.481 |
| | | RF | 0.481 | 0.424 | 0.384 | 0.474 | 0.450 |
| | Control | BLUP | 0.618 | 0.633 | 0.612 | 0.624 | 0.618 |
| | | RF | 0.624 | 0.640 | 0.614 | 0.594 | 0.588 |
| TillerLength_cm | Drought | BLUP | 0.343 | 0.383 | 0.351 | 0.483 | 0.389 |
| | | RF | 0.470 | 0.493 | 0.482 | 0.475 | 0.510 |
| | Control | BLUP | 0.573 | 0.600 | 0.560 | 0.615 | 0.542 |
| | | RF | 0.562 | 0.573 | 0.586 | 0.558 | 0.556 |
| **Sum (Mean)** | Drought | BLUP | 0.440 | 0.424 | 0.441 | 0.464 | 0.452 |
| | | RF | 0.424 | 0.411 | 0.408 | 0.415 | 0.437 |
| | Control | BLUP | 0.580 | 0.608 | 0.584 | 0.582 | 0.567 |
| | | RF | 0.568 | 0.591 | 0.595 | 0.554 | 0.559 |

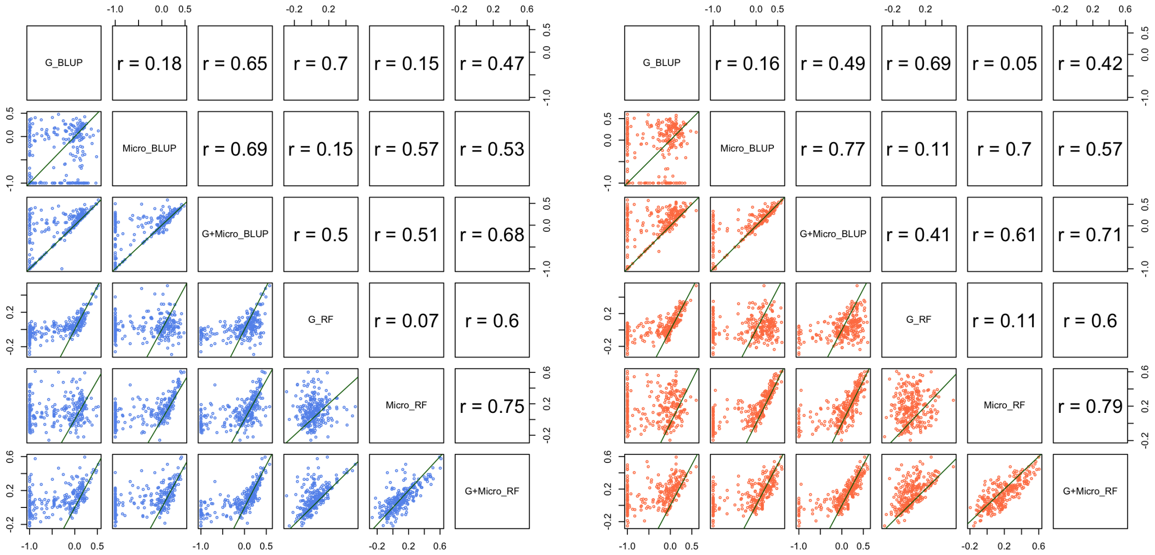


Figure 5S: Meta-metabolome prediction; the pair plot of the correlation coefficient of the model fitting, intra-year model (tested and trained in 2019). Each dot represents one metabolite.

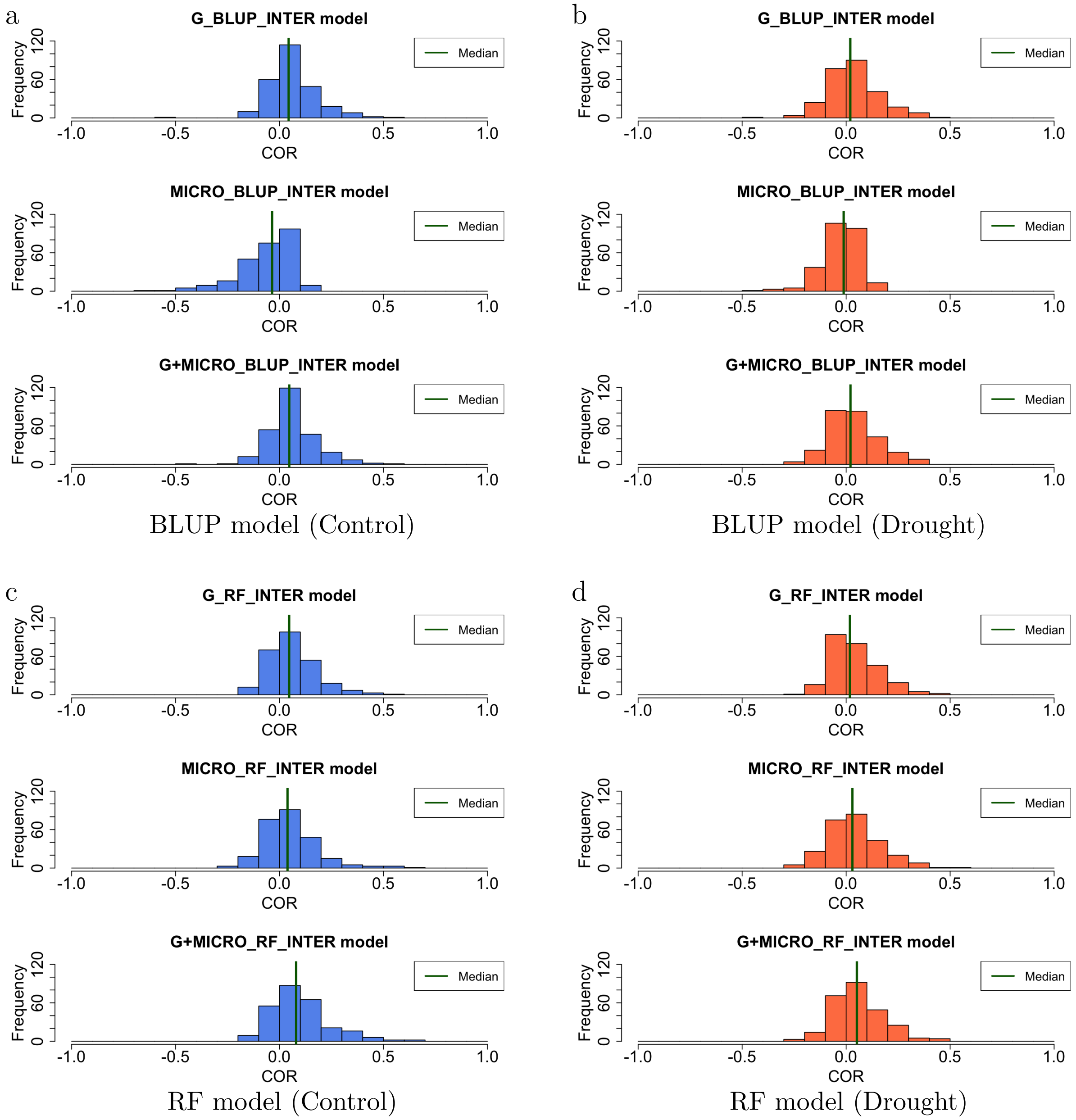


Figure 6S: Metabolite Prediction in the 2019 Drought Inter model (Model trained in 2020): Histogram of model accuracy of each metabolite by BLUP (Left) and by RF (Right). The input of the model is Genome (Top), Microbiome (Middle), both Genome and Microbiome (Bottom). The green bar represents the median.

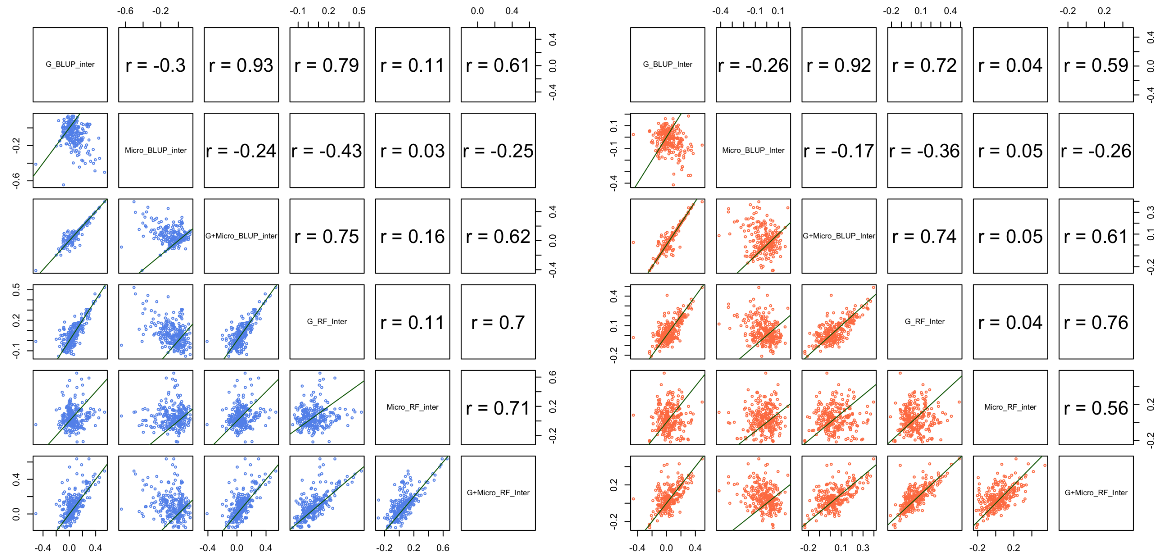


Figure 7S: Meta-metabolome prediction; the pair plot of the correlation coefficient of the model fitting, inter-year model (Model trained in 2020 and tested in 2019). Each dot represents one metabolite

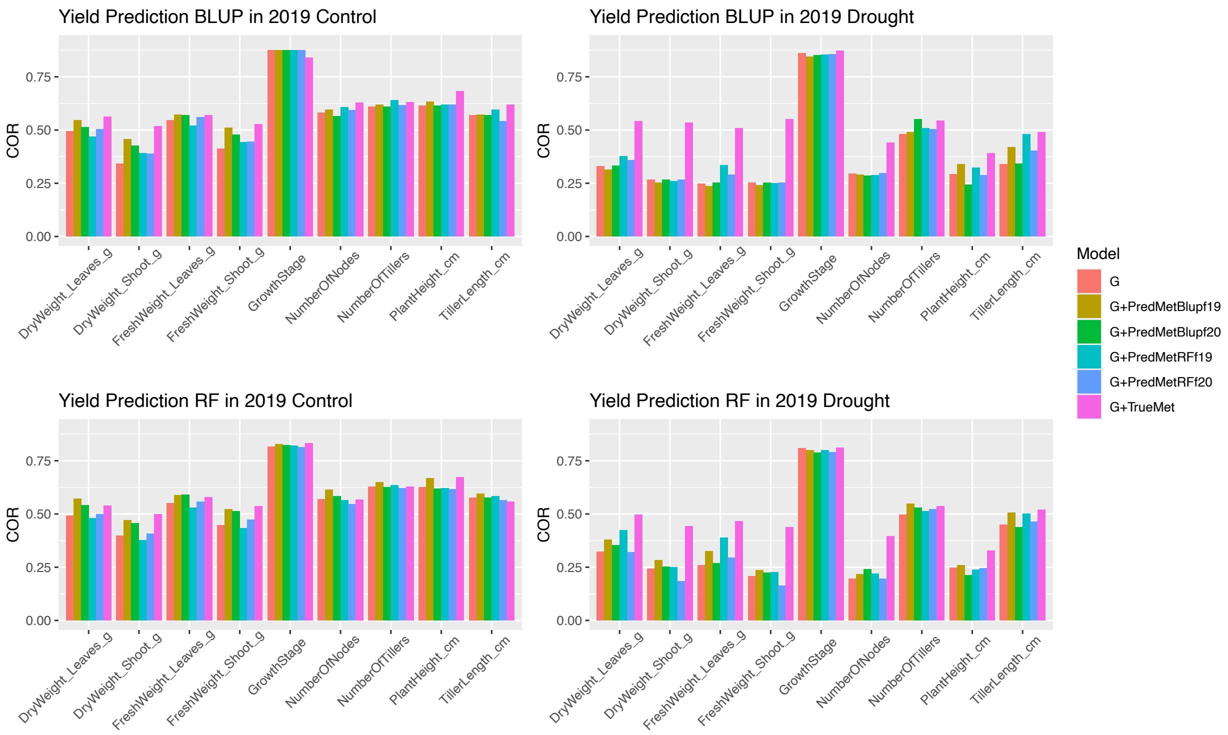


Figure 8S: Phenotype Prediction in the 2019 G-Proxy model: The Y-axis shows the correlation coefficient of the model fit to each phenotype. The phenotype prediction model is based on BLUP and tested with different inputs, such as G; genome, TrueMet; the actual meta-metabolome data in 2019, PredMet*ModelfYY*; "proxy" meta-metabolome predicted by *Model* in *YY* with Model= (BLUP, RF), YY=(19,20). "+" indicates that the model has multiple inputs.

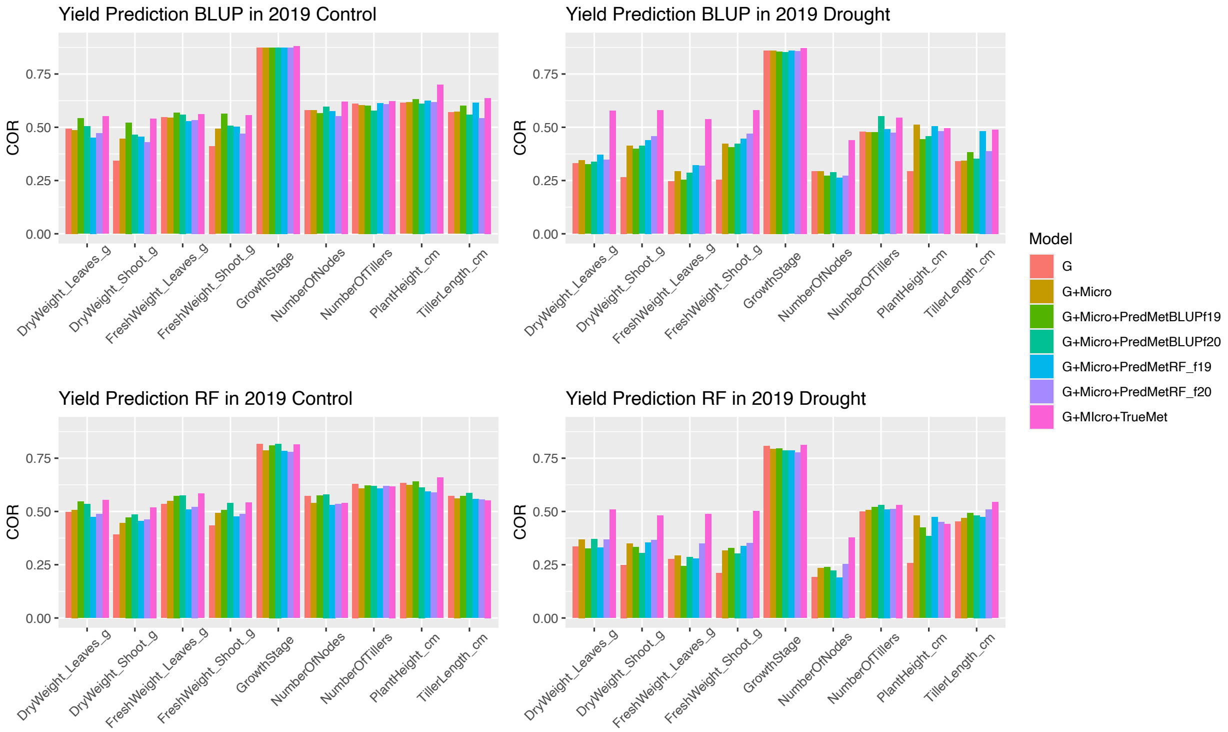


Figure 9S: Phenotype Prediction in the 2019 G-Micro-Proxy model: The Y-axis shows the correlation coefficient of the model fit to each phenotype. The phenotype prediction model is based on BLUP and tested with different inputs, such as G; genome, Micro; microbiome, TrueMet; the actual meta-metabolome data in 2019, PredMet*ModelfYY*; "proxy" meta-metabolome predicted by *Model* in *YY* with Model= (BLUP, RF), YY=(19,20). "+" indicates that the model has multiple inputs.

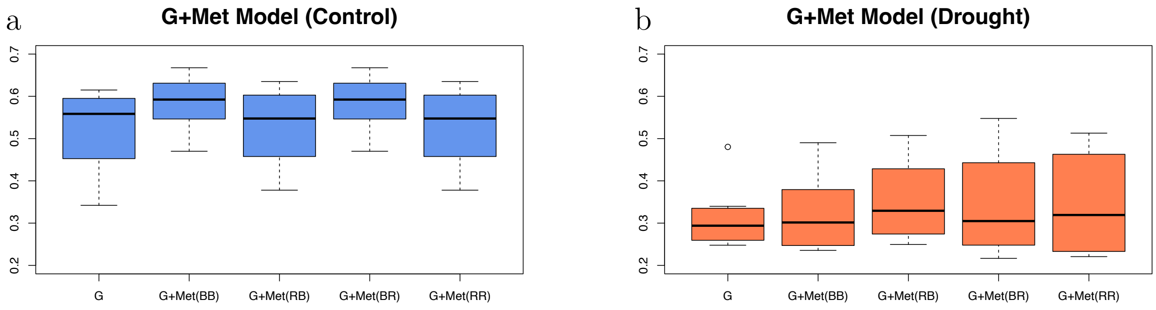


Figure 10S: Model selection of G-Proxy Model. B and R represent the model selection in each step. If BB, it's BLUP-BLUP model, RR is RF-RF model. In control, the best model is BB model, while the best model is RF-BLUP for drought. For this figure, the Growth Stage is excluded as an outlier.

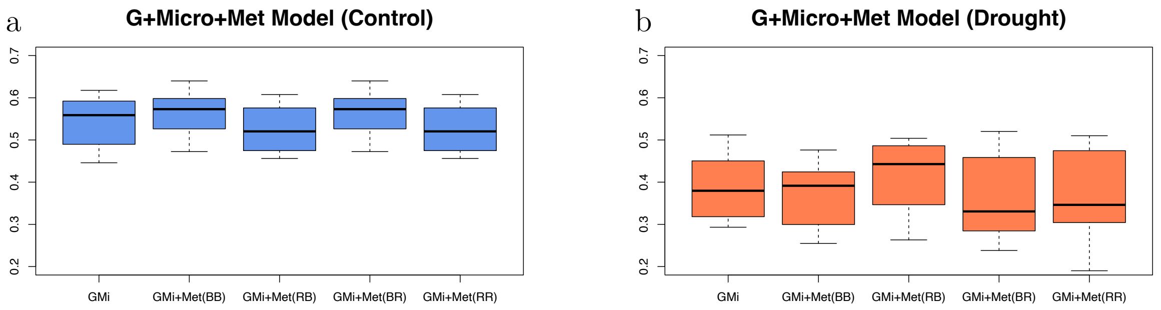


Figure 11S: Model selection of G-Micro-Proxy model. B and R represent the model selection in each step. If BB, it's BLUP-BLUP model, RR is RF-RF model. In control, the best model is BB model, while the best model is RF-BLUP for drought. For this figure, the Growth Stage is excluded as an outlier.

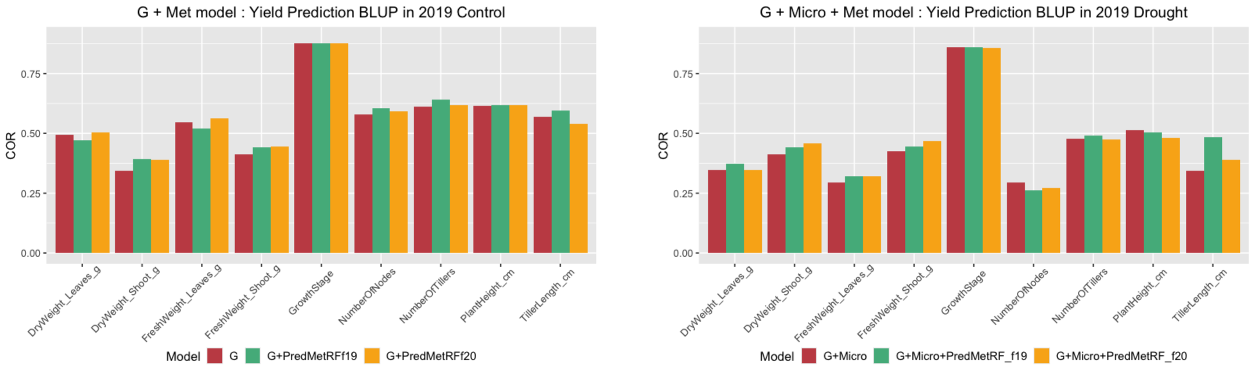


Figure 12S: Phenotype Predictions. The G-Proxy models were used for phenotype predictions, where G represents Genome, Met represents Meta-metabolome. The first step involved predicting intermediate meta-metabolomic traits using RF, followed by BLUP for the succeeding phenotype prediction. The legend shows whether the model was trained within the same year (19: Intra-year) or across years (20: inter-year). Each data point represents a phenotypic trait. Each model was tested using LOOCV, with correlation coefficients calculated and standardized.

### 1.2 Field experiment

The accessions and experimental fields were the same as those utilized by Toda et al. (2022) and Sakurai et al. (2022). A diverse collection of 198 soybean accessions, registered with the NARO Gen-bank (`https://www.gene.affrc.go.jp/`), was employed. This group primarily comprises global soybean minicore collections (Kaga et al., 2012; Kajiya-Kanegae et al. 2021).The field trial took place in 2019 and 2020 at the Arid Land Research Center of Tottori University on sandy soil (35°32' N, 134°12' E, 14 m above sea level). Each plot contained four plants, spaced 50 cm between rows, 80 cm between plots, and 20 cm between individuals. Sowing took place in early July, followed by thinning two weeks later. Fertilizer (13, 6.0, 20, 11, and 7.0 g m-2 of N, P, K, Mg, and Ca, respectively) was applied before sowing. White mulch sheets (Dupont, Wilmington, DE, USA) were laid to prevent rainwater infiltration and manage soil conditions by artificial irrigation. Watering tubes were placed under the leaves for field irrigation. Two watering treatments, no watering and watering, were applied to assess the effects of drought and control conditions, beginning after thinning and two weeks after sowing. Artificial irrigation was administered at a rate of 1.1 L/h per $m^2$ for 5 hours per day (7:00–9:00, 12:00–14:00, and 16:00–17:00). For phenotypic data sampling, the second and third individuals were sampled in each plot. Root sampling took place in early September and was performed on the third individual. A root sample was collected and stored at -80 degrees Celsius for subsequent meta-metabolome and microbiome analysis. For the details of meta-metabolome and microbiome analysis, refer to Dang et al. (2023, bioRxiv). For the details of the protocols of meta-metabolome analysis, also refer to Sawada et al. (2009) and Uchida et al. (2020). For the details of the protocols of microbiome analysis, refer to Kumaishi et al. (2022) and Ichihashi et al. (2020). The metabolome data were downloaded from the RIKEN DropMet website (`http://prime.psc.riken.jp/menta.cgi/prime/drop_index`; ID: DM0071, DM0072).

### 1.3 16S V4 rRNA data preprocessing

The raw paired-end sequencing reads were processed using QIIME2 (version 2020.6.0). The Raw FASTQ files were imported into QIIME2, and sequencing primers were removed using the Cutadapt plugin. For amplicon sequence variant (ASV)-based analysis, the primer-free reads were processed through the DADA2 pipeline, as described by Callahan et al. (2016), which included truncating the forward reads to 240 bp and the reverse reads to 180 bp, followed by filtering, denoising, merging, and chimera removal. Taxonomy assignment for all ASVs was performed using the SILVA reference database (version 138) (Quast et al. 2012; Yilmaz et al. 2014). Sequences classified as Archaea, Eukaryota, mitochondria, or chloroplasts were excluded from further analysis.

### 1.4 Supplementary equations

#### Graphical equations

The proof of (5) is as follows

$$
\begin{aligned}
P(B_{\text{pred}} \mid A, D) &= \frac{P(B_{\text{pred}}, A, D)}{P(A, D)} \\
&= \frac{P(B_{\text{pred}} \mid A)P(A)P(D)}{P(A)P(D)} \\
&= P(B_{\text{pred}} \mid A)
\end{aligned}
$$

Here, we assume that $A$ and $D$ are independent, so that

$$P(A, D) = P(A)P(D)$$

### Nonlinearity from A to C and A to B

Following the nonlinear scenarios (9), (10), and (11), we can introduce nonlinearity to the connection between $A$ and $C$ as follows.

$$C_{relu} = ReLU(A)\theta_{ac} + B_{relu}\theta_{bc} + D\theta_{dc} + \epsilon_c \tag{12}$$

$$C_{square} = A^{\circ 2}\theta_{ac} + B_{square}\theta_{bc} + D\theta_{dc} + \epsilon_c \tag{13}$$

$$C_{sigmoid} = \sigma(A)\theta_{ac} + B_{sigmoid}\theta_{bc} + D\theta_{dc} + \epsilon_c \tag{14}$$

These terms, $C_{\text{relu}}, C_{\text{square}}$, and $C_{\text{sigmoid}}$, incorporate nonlinearity in the data generation process. The parameters were aligned with those used in the proposed the linear and the nonlinear scenarios.
